# A single cell ATAC-seq atlas uncovers dynamic changes in chromatin accessibility during cell fate specification at the neural plate border

**DOI:** 10.1101/2025.05.03.652017

**Authors:** Eva Hamrud, Jacob Leese, Alexandre P Thiery, Ailin L. Buzzi, Alessandra Vigilante, James Briscoe, Andrea Streit

## Abstract

During development, dynamic changes in gene expression and chromatin architecture drive the transition from progenitors to specialised cell types. Here we use single cell ATAC sequencing (scATAC-seq) to investigate changes in chromatin accessibility as neural plate border cells segregate into neural, neural crest and placode cells. We developed a Nextflow pipeline, ‘**s**ingle **c**ell **A**dvanced **C**hromatin **E**xploration’ (scACE), which integrates scATAC-seq and scRNA-seq data to identify cell state specific accessibility profiles and groups of chromatin regions with coordinated dynamic behaviour, termed accessibility modules (AMs). We find that progenitors are characterised by broadly open chromatin, reflecting their broad potential to generate any ectodermal derivative. As development proceeds, cell-type-specific chromatin signatures are established. Inferring an enhancer-centric gene regulatory network, we predict Foxk2 as new regulator for placode specification and verify this prediction experimentally. Foxk2 target enhancers are open in placodal, but not any other ectodermal cells. This finding suggests that on a regulatory level, cells can use different strategies to control fate choice: differential accessibility of enhancers and broad accessibility controlled by differentially expressed transcription factors.

## Introduction

During embryonic development, a sequence of instructions directs cells along different pathways towards distinct identities. This directional process entails a series of decision events driven by changes in gene expression, which in turn is controlled by the interplay between transcription factors and non-coding cis-regulatory elements (CREs) that integrate both cell intrinsic and extrinsic information. Together, their dynamic changes over time form the gene regulatory networks (GRNs) that ultimately lead to the stabilisation of cell type specific genetic programmes^1^. Transcriptional profiling of single cells has revealed that individual progenitors initially co-express not only a handful of genes, but whole gene modules that are later specific for definitive cell types^2-4^. How such conflicting gene expression is resolved remains poorly understood, largely because regulatory information is often missing.

The vertebrate nervous system is complex both in its anatomy and cellular diversity. Yet, during development it arises from only three cell populations. The central nervous system originates from the neural plate, while the peripheral nervous system comes from two progenitor populations: neural crest cells and sensory placodes^5-8^. Among other cell types, neural crest cells generate neurons and glia along the entire body axis, while placodes are confined to the head and make major contributions to the sense organs and cranial sensory ganglia. As the neural plate forms, progenitors for all three cell types are located at its border and gradually diverge to acquire their unique identity. Recent single cell transcriptomic profiling ^4,9,10^ has revealed that neural plate border (NPB) cells cannot be defined by a unique transcriptional profile but co-express sets of genes that are later confined to the central nervous system, neural crest or placodes^4^. This observation suggests that their fate is ‘undecided’ and that they retain the ability to contribute to all three lineages even at relatively late stages. None of these studies, however, identified new transcription factors or target CREs that drive cells to adopt specific fates.

Here we use single cell Assay for Transposable-Accessible Chromatin^11^ (scATAC-seq) as a proxy to define CREs, and generate a high-resolution single cell chromatin accessibility atlas of the chick NPB and its derivatives from head process to the 8-somite stage (ss). To allow maximum accessibility, transparency and reproducibility of our data we have generated a user-friendly Nextflow pipeline, scACE (single cell Advanced Chromatin Exploration) which enables data exploration in a systematic, unbiassed and reproducible way (Figure 1) and integrates the new scATACseq data with previously published scRNAseq from equivalent cell populations^4^, effectively creating a pseudo-multiome data set.

**Figure 1.**
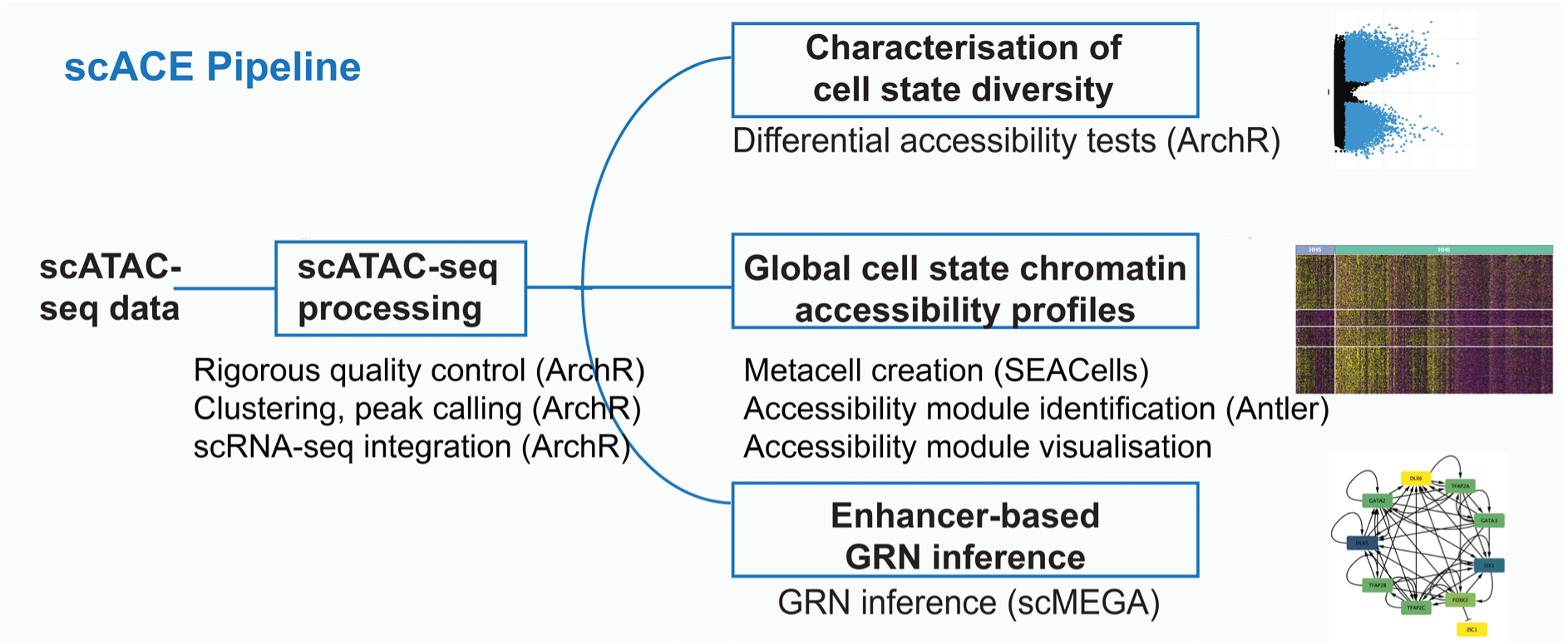
Nextflow pipeline scACE. After standard processing including quality control, clustering and peak calling scATAC-seq data points are integrated with equivalent scRNA-seq data. Processed and integrated scATAC-seq data are then analysed using three complementary approaches. 1) Differential accessibility tests explore how cell states diverge over time. 2) Accessibility modules (AMs) are generated and visualised to uncover global patterns in CRE activity over time. AMs are generated on metacells rather than single cells to overcome sparsity issues. 3) An enhancer-based GRN is inferred to identify new regulators of cell fate decisions. scACE Nextflow pipeline is available at https://github.com/Streit-lab/scACE.

We find that NPB cells are characterised by broad chromatin accessibility reflecting their competence to contribute to all ectodermal lineages. At early somite stages (ss), we identify accessibility patterns specific to placodal and neural cells, however, neural crest cells are instead characterized by profiles shared with both cell populations suggesting that they remain ‘undecided’ with the potential to contribute to more than one lineage until at least 8ss. To explore the mechanism by which stable ectodermal profiles emerge from the NPB, we inferred an enhancer-centric GRN, because integrating regulatory information considerably improves predictive accuracy and can be used to generate new testable hypotheses^12,13^. This approach identifies Foxk2 as a new regulator of *Six1*, a key gene required for placode identity^14,15^. Foxk2 is broadly expressed in the chick ectoderm but acts specifically in placodal cells because its target CREs are inaccessible in other cell types. This highlights differential accessibility as a mechanism for making fate decisions in which competence to respond to a ubiquitous transcription factor is restricted to a specific cell population.

## Results

### A single-cell chromatin accessibility atlas for the neural plate border

The neural plate border (NPB) contributes to the entire vertebrate nervous system and skin via four transient cell populations: neural plate, neural crest, placodal and future epidermal cells^6-8,16^. We and others have previously shown how gene expression in single cells gradually diverges as NPB cells adopt their distinct identities^4,9,10^. However, the corresponding CREs controlling these changes in gene expression have not been characterised at a single cell level. Here, we present a high-quality atlas of chromatin accessibility profiles NPB cells and their derivatives, with single cell resolution.

To measure chromatin accessibility at a single cell level, we performed 10 x Genomics single-cell ATAC sequencing (scATAC-seq) of chick NPB and adjacent cells at Hamburger and Hamilton^17^ (HH) stages HH5, HH6, HH7, HH8 (ss4) and HH9 (ss8) spanning the time period when cell fate decisions occur (Figure 2A). 85,396 high-quality nuclei were retained after quality control to account for differences in sequencing depth between samples (Supplementary Figure 1A-C). Using Uniform Manifold Approximation and Projection (UMAP) for dimensionality reduction and visualisation we identify 13 clusters (Figure 2C, Supplementary Figure 1D). Overlaying the UMAP with developmental stage reveals that global chromatin accessibility profiles gradually change from HH5 to ss8 (Figure 2B). Most clusters contain cells from multiple stages (compare Figure 2B and C) and we confirm this plotting heatmaps showing the relative proportion of cells in each cluster and each stage (Supplementary Figure 1E). Thus, although there is a gradual shift in chromatin accessibility over time, cells from successive developmental stages share similar chromatin accessibility profiles.

**Figure 2.**
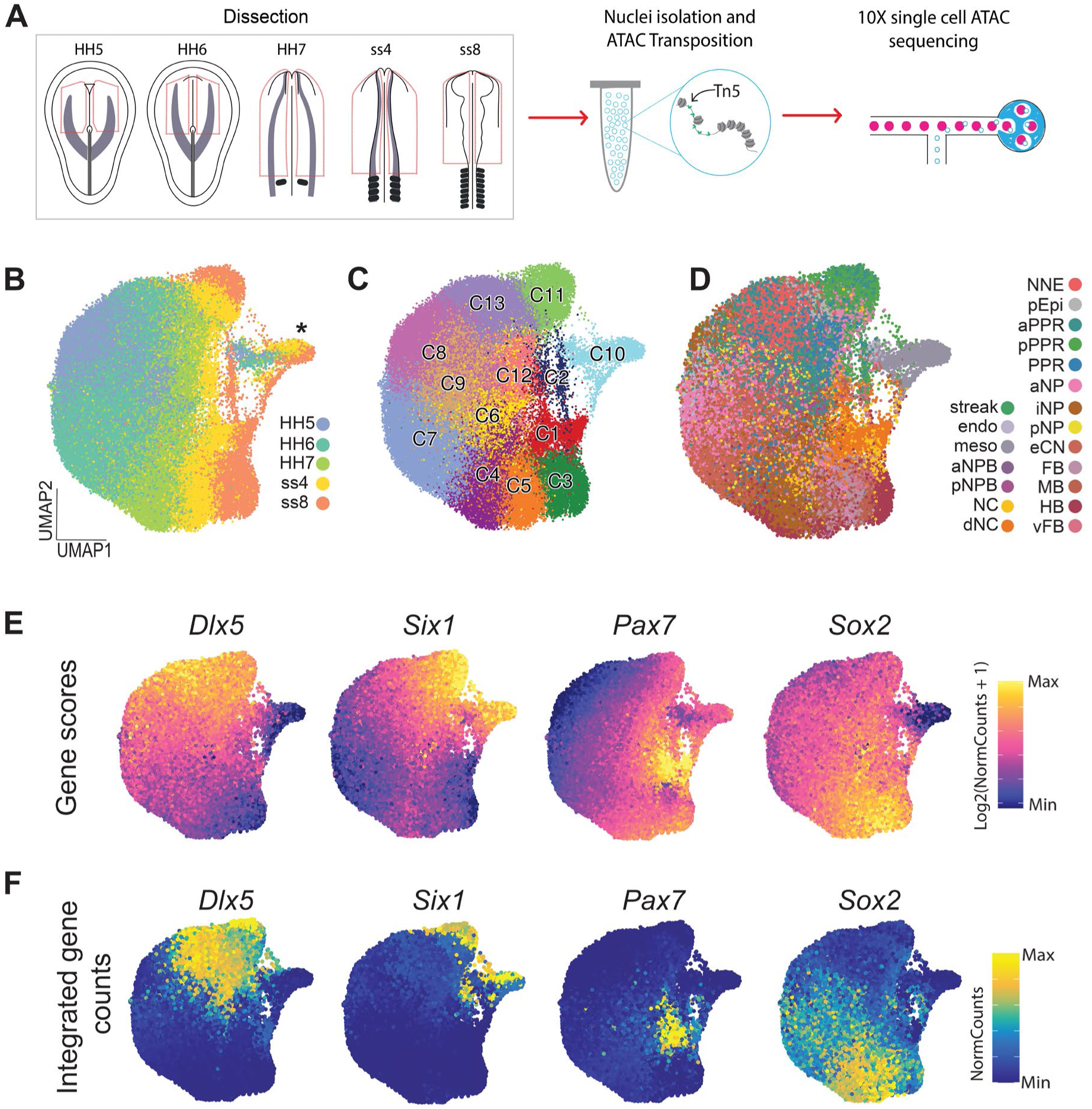
A chromatin accessibility atlas of the developing chick ectoderm. A. Workflow of scATAC-seq data collection. Regions outlined in red were dissected from chick embryos at five developmental stages (HH5, HH6, HH7, ss4 and ss8). Single nuclei were isolated and processed for 10x scATAC sequencing. 15,929 nuclei were from HH5, 33,189 from HH6, 16,196 from HH7, 11,979 from ss4 and 8,102 from ss8. B. UMAP embedding of scATAC-seq atlas overlaid with developmental stage information. * labels cluster C10 composed of cells from many developmental stages C. UMAP embedding of scATAC-seq atlas overlaid with cluster assignments. D. UMAP embedding of scATAC-seq atlas overlaid with cell state annotations transferred from published equivalent scRNA-seq data^4^. E. Gene score feature plots for key genes as inferred from scATAC-seq data. F. Gene expression feature plots for key genes as transferred from scRNA-seq data.

To annotate cell states we integrated the scATAC-seq data with previously published scRNA-seq data from equivalent stages^4^ using the scACE pipeline. Using promoter accessibility as a proxy for gene expression we paired scATAC-seq and scRNA-seq cells based on similarity of gene expression, and transferred the cell identity label ^18^. Paired cells are distributed evenly across cell states and stages (Supplementary Figure 2A) and the proportions of cells assigned to different identities are almost identical in both data sets (Supplementary Figure 2B, C). To confirm the accuracy of our data integration, we compared gene expression measured in the scRNA-seq data with gene expression inferred from promoter accessibility. We find that the expression levels of key markers from both modalities closely match and display expected patterns (Figure 2E, F). For example, *Six1* is enriched in placodal cells (C11), *Dlx5* expression spans both placodal and non-neural ectoderm (C8, C11, C13), *Pax7* is restricted to cells labelled as neural crest (C1) and *Sox2* to neural cells (C5, C3).

To identify putative enhancer regions from the scATAC-seq data we called peaks using ArchR’s implementation of MACS2^19^. Of 268,323 peaks called across all five developmental stages most are located in intronic and distal genomic regions and may therefore represent active CREs (Supplementary Figure 3A). To explore this possibility, we cross-referenced our scATAC-seq atlas with previously characterised chick CREs known to have spatially restricted activity at ss8 (Supplementary table 1). Of 68 published enhancers examined, 21 maintained similar accessibility across all ectodermal cell populations, while the remaining 47 showed differential accessibility patterns consistent with their reported *in vivo* activity. For example, the Sox10_E1 enhancer, which is active in the neural folds and migrating neural crest cells^20^, shows highly specific accessibility in neural crest cells (Supplementary Figure 3B), while the Six1-21 enhancer, which is active in the olfactory, otic and epibranchial placodes^21^, shows increased accessibility in placodal cells (Supplementary Figure 3C; pre-placodal region [PPR], anterior and posterior PPR).

In summary, we present a single cell chromatin accessibility atlas of the chick NPB and its derivatives. Each data point contains accessibility as well as gene expression data, effectively creating a pseudo-multiome dataset for further exploration.

### Characterising changes of chromatin accessibility

Sampling single cells across developmental time allows us to assess how chromatin accessibility changes as progenitors generate definitive neural crest, placodal and neural cells. To investigate this, we generated separate UMAPs for each developmental stage independently. At HH5 and HH6 most cells do not segregate based on chromatin accessibility but rather form a diffuse cloud without distinct clusters (Figure 3A). Most clusters contain multiple intermixed cell states (Figure 3A, B, Supplementary Figure 4A, B). By HH7, cells are broadly organised into neural (clusters C4, C7-9) versus placodal (cluster C3) identities, although three clusters (C1, C5, C6) continue to encompass a mixture of both cell states (Figure 3A, B, Supplementary Figure 4C). Although we can identify neural crest cells at HH7, they are widely distributed across clusters and only by ss4 localise to a distinct cluster. Thus, neural crest cells lack a unique chromatin accessibility profile before early somite stages. In addition, several clusters contain a mixture of cells even at ss4 and ss8 (Figure 3A, B; Supplementary Figure 4D, E). C7/8 encompass placode and anterior neural plate cells, C3 hindbrain and neural crest cells and C4 all three cell types. To confirm that chromatin accessibility profiles become more distinct as development proceeds, we used the scACE pipeline to perform differential accessibility tests between clusters at each stage. The number of differentially accessible peaks, as defined by an FDR of less than 0.05, dramatically increases over developmental time (Figure 3C).

**Figure 3.**
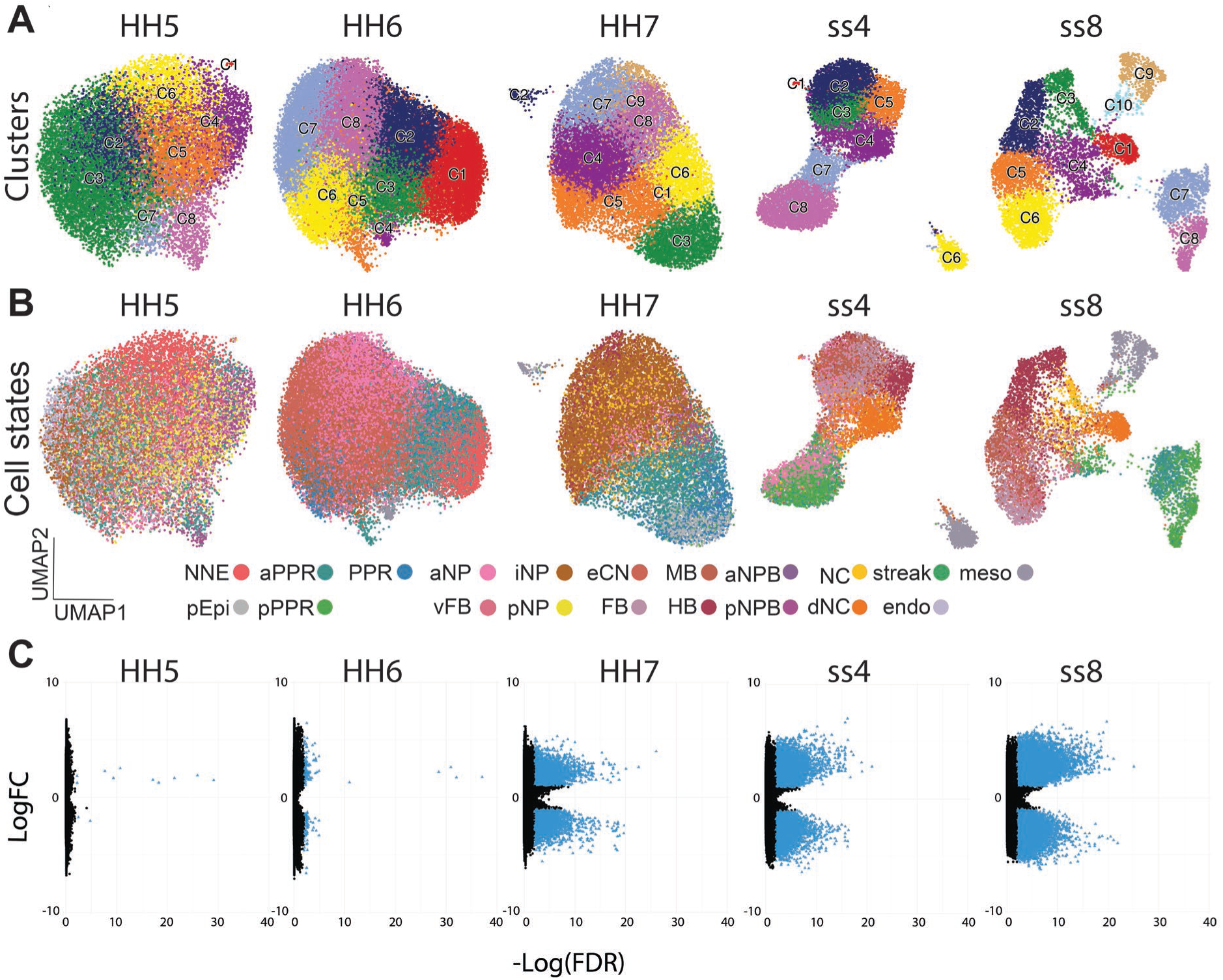
Cell state diversity during progenitor cell diversification. A. UMAP embeddings of scATAC-seq data for each developmental stage overlaid with cluster assignments. B. UMAP embeddings of scATAC-seq data for each developmental stage overlaid with cell state annotations. C. Volcano plots of differential accessibility tests between clusters at each developmental stage. Blue points indicate significant hits of FDR less than 0.05, black points indicate non-significant hits.

Together, these data show that NPB cells initially share chromatin accessibility profiles. As they diversify into neural, neural crest, placodal and future epidermal lineages their accessibility profiles diverge. Despite this segregation, some clusters continue to contain mixed cell states suggesting that their final fate remains undecided.

### Dynamic and coordinated changes in chromatin accessibility

Typically, characterisation of cell states in scATAC-seq data involves comparing clusters using differential accessibility tests. However, this approach fails to uncover shared accessibility profiles and their dynamics, which may in turn correspond to co-regulated enhancer modules that integrate cell fate decisions. To characterise chromatin regions which open and close in a coordinated manner as NPB progenitors acquire distinct identities, we clustered chromatin regions into groups called accessibility modules (AMs). As the sparsity of scATAC-seq precludes direct clustering of chromatin regions, we first grouped single cells into metacells, which provide a fine-grained view of the data and cleanly separate into cell states.

Metacell generation was performed using the SEACells package^22^ in the scACE pipeline. Each metacell contains on average 40 cells with closely matching accessibility profiles. We also calculated metacells from the equivalent scRNA-seq data^4^ to overcome potential issues of RNA dropouts. Each metacell is highly homogeneous: contributing cells have the same cell state, come from the same cluster and have highly similar accessibility or transcriptional profiles. At the same time, different metacells are well separated and thus capture the diversity of different cell states observed in the single cell data (Supplementary Figure 5). To annotate cell states of the ATAC metacells, both modalities – gene expression and accessibility - were reintegrated at the metacell level using the same approach as for single cells (Supplementary Figure 6).

Next, we calculated AMs by repurposing the Antler package originally created for scRNA-seq data analysis^23^. Amalgamating peak counts across metacells rather than single cells allows the calculation of peak-peak Spearman correlation. Considering only the top 10,000 most variable peaks in intronic or distal regions (i.e. potential CREs), we performed hierarchical clustering and defined 15 AMs across the full dataset (Supplementary Table 2). Each AM contains a group of chromatin regions that open and close together across all ATAC metacells. Of these, three modules were either globally open or closed at each timepoint and were not considered for further analysis.

Visualising the remaining 12 AMs using heatmaps reveals that 4 are open in placodal cells at ss8, while 8 are accessible in neural cells. Of the placodal modules, AM1 and AM4 show restricted pan-placodal accessibility, while AM2 and AM3 are also accessible in neural crest cells (Figure 4A, C). Neural AMs show clear regional patterns: AM6 and AM7 are preferentially open in hindbrain cells, while AM10 and AM11 are biased towards forebrain (Figure 4A, C). AMs 12-15 are open in most neural cells and in neural crest cells although AM 13-15 appear to close when neural crest cells delaminate (Figure 4A, C). Thus, while neural and placodal cells are characterised by unique AMs, neural crest cells share at least some putative enhancer regions with neural and placode cells.

**Figure 4.**
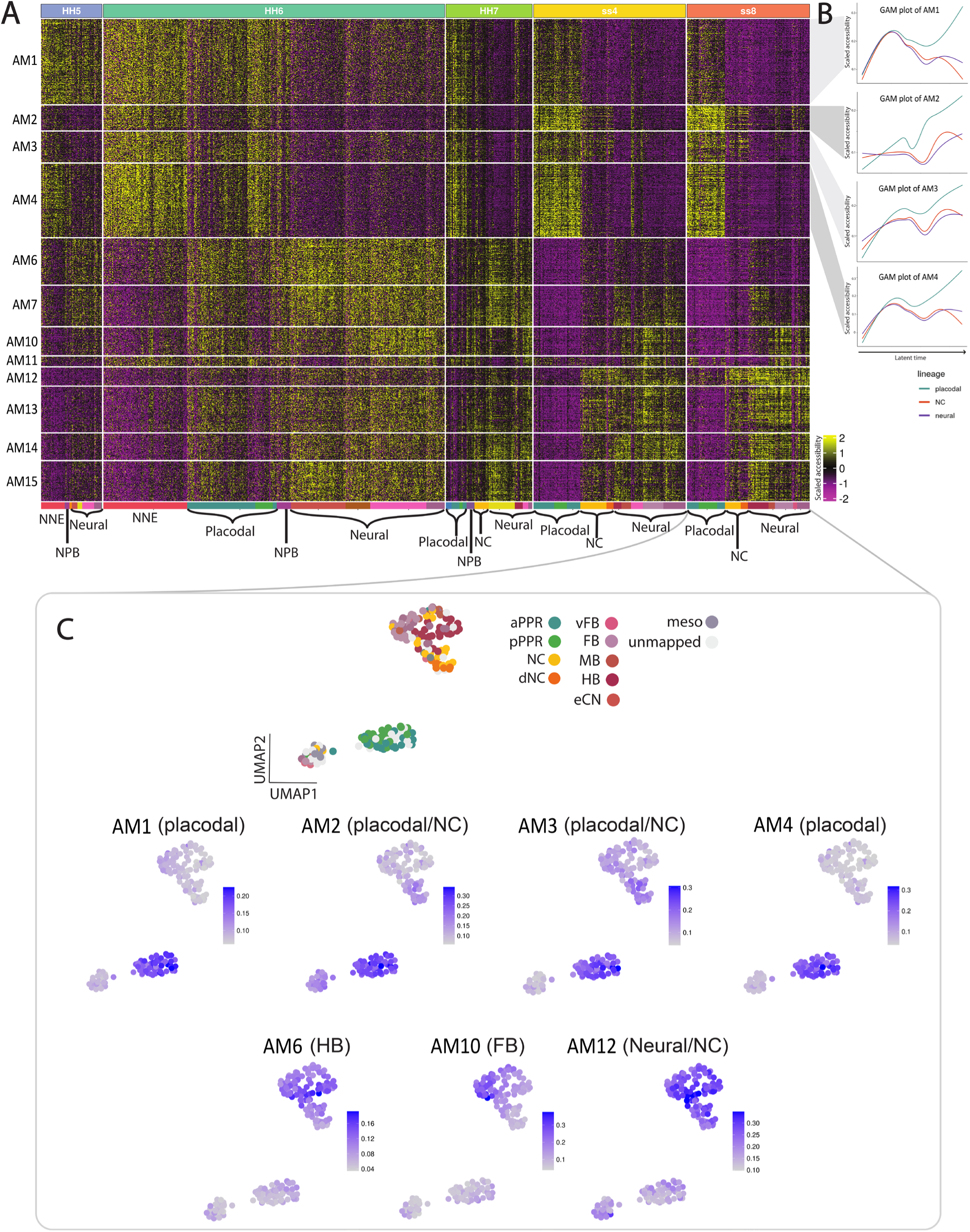
Global characterisation of chromatin accessibility profiles. A. Heatmap of 12 accessibility modules (AMs) showing cell-state specific accessibility. Scaled accessibility is shown per metacell, organised by developmental stage and within each stage by cell state. B. GAM plots showing scaled accessibility of placodal modules AM1, AM2, AM3 and AM4, weighted by neural, neural crest and placodal lineages. The latent time and lineage probabilities were reused from Thiery and colleagues^4^. C. Top: UMAP embedding of metacells at ss8 overlaid with cell state; below: feature plots of the average accessibility of each AM across metacells.

Assessing the dynamic changes of AMs over time reveals distinct behaviours of enhancer groups. At HH6, modules AM6-15 are accessible in all cells except those of the NNE. By HH7 these modules have largely closed in placodal cells and by ss8 become restricted to neural cells of various rostro-caudal identities (AM6-10) or to both neural and neural crest cells (AM12-15) (Figure 4A). Using generalised additive models (GAMs), we visualised the dynamic accessibility changes of AM1-4 across placodal, neural and neural crest lineage trajectories (inferred using RNA splicing kinetics^4^) (Figure 4A). Chromatin regions in AM1 and AM4 are broadly accessible in nearly all cells initially, then gradually close in neural and neural crest cells, to become confined to placodal cells by ss8. In contrast, regions in AM2 and AM3 show preferential accessibility in placodal, non-neural and NPB cells at HH6 and after HH7 remain accessible in placodal and neural crest cells (Figure 4A, B). Motif enrichment analysis of the open chromatin regions in each placodal module reveals a strong enrichment of Tfap2a motifs in AM3 and of both Tfap2a and Gata motifs in AM4 (Supplementary Fig. 7), suggesting that these transcription factors regulate the CREs within these modules. This finding is consistent with the established roles of these factors in specifying neural crest and placode cell fates^24-29^.

In summary, AM analysis implemented in the scACE pipeline allows unbiased characterisation of global changes in chromatin accessibility profiles. Grouping single cells into metacells overcomes data sparsity of scATAC-seq data while generating a fine-grained view of chromatin accessibility changes. We identified placode- and neural-specific AMs which may participate in driving the emergence of these fates. However, neural crest cells share accessibility with placodal or neural cells suggesting they remain plastic.

### NPB and neural crest cells are characterised by co-accessibility of neural and non-neural AMs

The analysis above reveals that NPB cells are not characterised by a unique accessibility profile but rather by co-accessibility of AMs that are later confined to specific cell populations (Figure 4A). This agrees with our previous finding that NPB cells co-express gene modules that are later specific for definitive neural, neural crest and placodal cells ^4^. Like NPB cells, NC cells display a mixed chromatin accessibility profile (Figure 4A). At ss8, they share accessibility patterns with placodal AMs 2 and 3 and neural AMs 12-15.

To examine co-accessibility in more detail, we used generated co-accessibility UMAPs displaying average AM accessibility across cells from each stage. This analysis reveals that the placodal AM3 and the neural AM12 are only co-accessible in a few metacells at early stages (HH5-7) but are co-accessible specifically in neural crest cells at ss4 and ss8 (Figure 5A, B). Comparing the placodal AM3 and all neural AMs (AM12-15) consistently shows their co-accessibility in NC cells (Supplementary Figure 8A-C).

**Figure 5.**
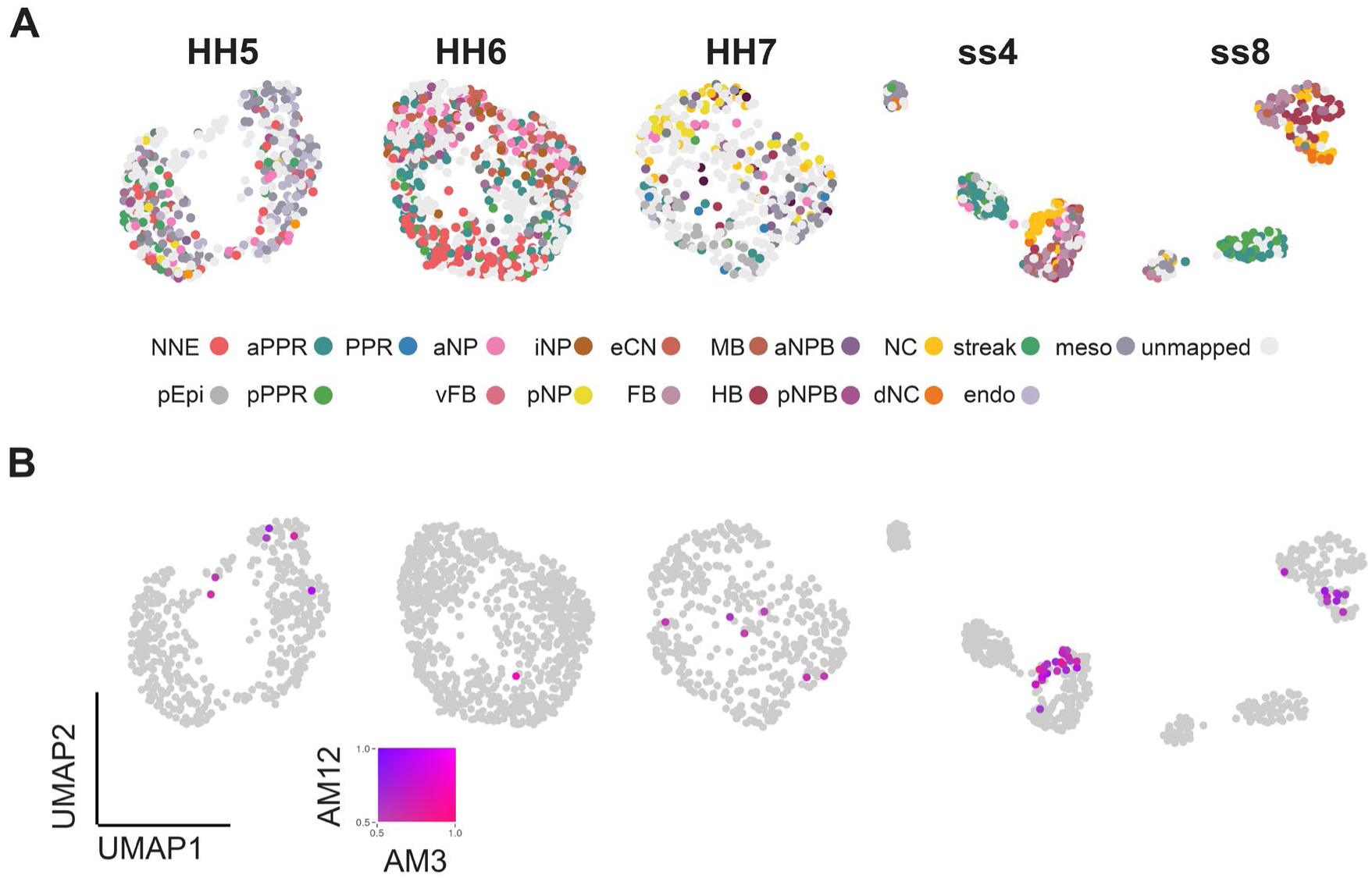
Neural crest cells are characterised by co-accessibility of neural and non-neural accessibility modules. A. UMAP embeddings of metacells at HH5, HH6, HH7, ss4 and ss8 overlaid with cell state annotations. B. Feature plots showing co-accessibility of the neural AM12 and the placodal AM3 at each developmental stage.

Together, these data suggest that, based on their chromatin accessibility profiles, NPB cells may have broad potential to give rise to all three fates. Among these progenitors, neural and placodal cells are the first to emerge, each with a unique accessibility profile, while neural crest cells display mixed profiles and therefore may retain broad potential at ss8. This implies that neural crest either remain ‘undecided’ even at later developmental stages or are specified by a mechanism distinct from that of placode and neural cells.

### Enhancer-centric GRN inference identifies new regulators of placode identity

To identify new regulators as NPB cells diversify, we leveraged our pseudo-multiome data to infer an enhancer centric GRN. Focusing on the placodal lineage, our GRN-inference approach uses an adapted version of the scMEGA package^30^. As potential regulators, we included all transcription factors that correspond to motifs present in accessible chromatin regions. We defined target nodes as genes that vary across the placodal trajectory using scRNAseq data^4^ (Figure 6A; 1,693 genes in total; Supplementary Table 3). Using ChromVar^31^, we then determined the activity of the transcription factors by measuring the accessibility of their target CREs. Regulatory interactions between these transcription factors and their target genes were inferred based on two criteria: i) the expression of the target gene must correlate with the accessibility of its associated enhancer containing the predicted transcription factor binding site and ii) the transcription factor activity must correlate with the expression of the target gene. To identify factors that promote placodal character, we only considered those that positively correlate with placodal genes resulting in the prediction of 24,058 direct regulatory interactions between 142 transcription factors and 1,431 targets (Figure 6A, Supplementary Table 3).

**Figure 6.**
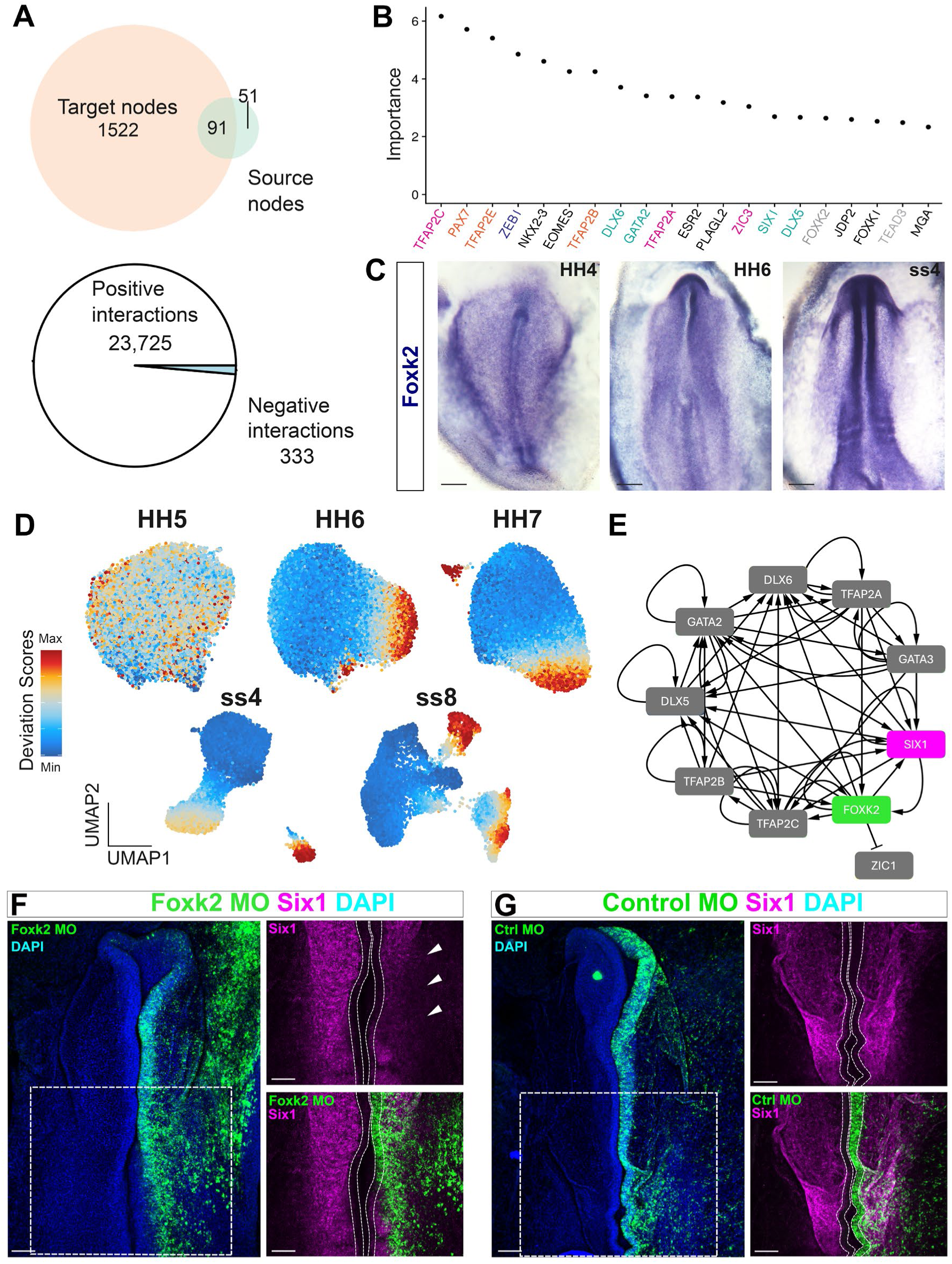
Enhancer-centric GRN inference identifies a new *Six1* regulator. A. Top: Venn diagram showing number of source and target nodes in the placodal GRN. Bottom: Pie chart showing the number of predicted regulatory interactions in the GRN. B. Plot of network analysis identifying top 20 most important factors based on betweenness and Page Rank scores. Factor names are coloured based on known activity: placodal and NC (pink), NC (orange), neural (blue), placodal (green). C. *In situ* hybridisation showing *Foxk2* expression. Scale bars: 200 μm. D. ChromVar feature plots of Foxk2 across developmental stages indicating Foxk2 activity. E. Network visualisation of the interactions between Foxk2 and key known placodal transcription factors. F. Foxk2 knock-down (green) leads to reduction of *Six1* expression (magenta); nuclei are labelled with DAPI (blue). Boxed area on the left is shown at higher magnification on the right. White lines indicate neural folds; white arrowheads: *Six1* reduction. Scale bar: 100 μm. G. Control morpholinos (green) do not affect *Six1* expression (magenta); nuclei are labelled with DAPI (blue). Boxed area on the left is shown at higher magnification on the right. White lines indicate neural folds. Scale bars: 100 μm.

To predict ‘hub’ nodes that transfer information across the network, we performed network ‘prioritisation’ analysis using scMEGA^30^. This analysis highlights several factors known to regulate placode fates as important hubs, including Six1, Dlx5 and -6, Gata and Tfap2 family members ^15,24-26,29,32-38^ (Figure 6B) adding confidence to the predictive power of this approach. In addition, other factors emerge that have not been implicated in placode precursor specification. Among those, Foxk2 is ranked highly in the network analysis (Figure 6B) and is predicted to regulate *Six1* expression directly (Figure 6E). Its target enhancers are uniquely accessible in the placodal/non-neural ectoderm but inaccessible in neural and neural crest cells (Figure 6D; compare to Figure 3B). In contrast, scRNA-seq^4^ shows that *Foxk2* mRNA is expressed broadly in the ectoderm, and we confirmed this here by *in situ* hybridisation (Figure 6C).

Thus, GRN inference predicts that Foxk2 specifically regulates placodal genes (i.e. *Six1)* but should not affect neural or neural crest markers, because its target enhancers are inaccessible in these cells. To test this, we knocked down *Foxk2* using splice-modifying morpholinos (MOs). We electroporated control or Foxk2-targeting MOs into one side of the chick epiblast at HH3/4 and assessed the expression of *Six1*, the neural marker *Sox2* and the neural crest marker *Pax7* at HH7-8 by *in situ* hybridisation chain reaction (HCR). We find that Foxk2 (n=7) but not control (n=5) MOs reduce or abolish *Six1* expression in electroporated cells (Figure 6F, G). In contrast, the expression of *Sox2* (n=8) and *Pax7* (n=10) is not affected in Foxk2 knock downs (Supplementary Figure 10).

In summary, we inferred an enhancer-centric GRN which contains over 20,000 regulatory interactions that may be active during the emergence of placodal cells. Network analysis recovers known regulators of placodal identity as well as new candidates, among them Foxk2. We experimentally verify a role for Foxk2 in controlling *Six1* expression and show that despite its broad expression it does not control the expression of neural and neural crest markers. Transcriptome analysis alone failed to identify Foxk2 as a new placodal regulator^4^, highlighting the power of multimodal approaches to discover new factors controlling cell fate choices.

## Discussion

The question of how cells of the embryonic ectoderm diversify to form progenitors of the central and peripheral nervous system has been extensively investigated. Characterisation of the underlying GRNs has revealed the signalling events that control the activation of neural, neural crest and placode specific gene expression as well as the transcriptional hierarchy as individual cell identities become distinct ^8,28,39-41^. However, the GRN structure was largely determined by experimental perturbation of one gene at a time and by assessing a handful of downstream targets, and with few exceptions, does not consider regulatory information.

Here, we generate a chromatin accessibility atlas of the chick NPB and its derivatives from head process to 8ss. We reveal how the chromatin landscape changes over time and define enhancer modules with coordinated dynamics that are likely to control fate decisions. Integrating the new scATACseq data with published scRNAseq data^4^ we generate a pseudo-multiome data set and use this to infer an enhancer-centric GRN. This analysis identifies Foxk2 as a new regulator of placodal identity and highlight chromatin remodelling events that open enhancers in specific cell populations as mechanisms to control cell fate choice and regulating the competence to respond to transcriptional regulators.

### Different mechanisms to control cell fate choice at the neural plate border

The NPB has been characterised as an ectodermal territory where neural and non-neural gene expression overlaps and where progenitors for the neural plate, neural crest, placodes and epidermis reside^5,7,8,42,43^. On the level of gene expression^4,10^ and chromatin accessibility (this study), NBP cells appear to be multipotent and do not show any bias towards a particular fate. Our findings point to different molecular mechanisms that control cell fate choice at the NPB.

On one hand, we reveal differential enhancer accessibility as a mechanism to control cell fate allocation at the NPB. This mechanism implies that cells are already primed for a specific decision with chromatin remodelling events opening CREs in a cell type specific manner and that the activating transcription factor(s) play a permissive rather than instructive role. A similar strategy is also implemented during ventral progenitor specification in the neural tube ^44^. Here, the pioneer factor Foxa2 plays a critical role in initiating chromatin changes. How placode specific Foxk2 target enhancers are made accessible in the placodal lineage is currently unknown.

On the other hand, we find that AMs which are open specifically in definitive neural, neural crest and placode cells at later stages, are co-accessible in NBP cells. This observation suggests that NBP cells are initially competent to execute any ectodermal programme, and that subsequently either differential binding of transcription factors activates cell type specific gene expression, or that accessible enhancers are gradually decommissioned by chromatin remodelling, or a combination of both. NPB cell plasticity is also evident by their co-expression of opposing gene modules including fate determinants like the placode specifier *Six1* and neural crest specifier *Pax7*^4,45^ raising the question of how such co-expression is resolved. Recent studies examining lineage determination in mouse blastocysts suggest that opposing fate determinants initially co-bind cell type specific enhancers, followed by rapid eviction of the ‘wrong’ transcription factor and rapid chromatin remodelling^46,47^. Whether similar mechanisms are at work during fate choice at the NBP remains to be elucidated.

### Emergence of different cell fates from the NPB

An open question is in which order neural, neural crest and placode cells emerge from NPB cells. Our analysis identifies groups of putative enhancers that open and close together which we term ‘accessibility modules’ allowing us to predict the order of events as different lineages are specified. We find that many chromatin regions are broadly accessible at early developmental stages but become later confined to placode precursors. This is reminiscent of changes in gene expression^4^: many genes that are expressed broadly at early stages become later restricted to placode cells. Together, these findings suggest that placodal identity may be a ‘ground state’ that must be repressed for other lineages to emerge. As placodal modules become restricted, neural-specific accessibility modules open around the time when neural-specific gene expression occurs^4^ indicating that neural cells emerge next.

These data are consistent with the idea that cells emerge from the NPB in a particular order. However, recent transcriptional profiling in Xenopus suggests a different scenario. Trajectory analysis using scRNAseq data of posterior NPB cells predicts that different routes can lead to the same fate^9^. For example, neural crest cells are proposed to arise either from partially neutralized ectoderm or directly from NPB cells accompanied by the expression of common and distinct genes. Likewise, two different paths may lead to placode precursors. This notion agrees with a recent model proposed in chick in which depending on their location cells have a different probability to generate definitive neural plate, neural crest and placodes^4^. They may do so taking different routes. Together, these observations highlight the complexity of fate decisions at the NBP and that such decisions take place over a protracted period. Ultimately, these predictions will need to be tested by in vivo lineage tracing.

### Plasticity of neural crest cells

Our analysis defines groups of enhancers open and close in a coordinated manner termed accessibility modules (AMs). While placodal and neural cells are characterised by unique AMs at ss8, neural crest cells are not. Instead, they share accessibility profiles with neural plate and placodal cells. This observation suggests that neural crest cells retain plasticity longer than neural and placodal cells in agreement with other findings. In chick pluripotency markers are expressed widely in the entire ectoderm^48,49^, however, the pluripotency factors Oct4 and Sox2 are repurposed specifically in neural crest cells to remodel the chromatin landscape^50^. In Xenopus, neural crest cells, unlike other ectodermal cells, maintain the expression of pluripotency markers and can not only generate ectodermal derivatives, but also endoderm and mesoderm when exposed to appropriate conditions^51^.

### Enhancer-centric GRN inference predicts a new regulator of placodal fate

The transcription factor Six1 and its cofactors Eya1 or Eya2 play a key role in placode precursor specification and subsequent placode development^14,15,33,52-58^. Members of the TFAP2, Gata and Foxi family are known upstream regulators of *Six1* and *Eya1/2*^25,28,29,59^. We find that TFAP2 and Gata motifs are highly enriched in AMs associated with placodal cells, suggesting that they may interact directly with placode specific CREs. Dlx and Msx transcription factors bind to a placode-specific *Six1* enhancer to activate or repress its expression, respectively^60^. However, beyond these findings, no direct regulators of the Six/Eya network have been identified. Our GRN inference approach identifies such regulators of *Six1* and placodal fate.

While many inference algorithms predict regulatory interactions based on the presence of transcription factor binding sites in CREs and differential expression of the relevant factor^61-63^, we used a modified version of scMEGA^30^ to infer a GRN, which does not consider differential expression of the regulating transcription factor. This analysis returns known *Six1* regulators as key network nodes but also highlights new factors including Foxk2. Modelling Foxk2 activity along the placodal trajectory reveals that its target enhancers are specifically active in placode precursors. In contrast, it is widely expressed in the early epiblast including the neural plate, NPB and future epidermis, and has therefore been overlooked in previous studies focussing on differential gene expression. As predicted, our functional experiments show that Foxk2 is specifically required for the expression of *Six1* in placode precursors but does not control neural and neural crest genes.

The role of Foxk2 during embryo development has not been investigated. Molecularly, it recruits repressor complexes to open chromatin shutting down gene expression^64-66^. However, more recently Foxk2 has also been implicated in transcriptional activation. In human embryonic stem cells, Foxk2 is largely associated with active histone marks and seems to ‘pre-mark’ CREs that are strongly activated as cells differentiate^67^. It is therefore possible that Foxk2 occupies placode-specific CREs that control genes required for placodal fate. Interestingly, Foxk2 is required for the recruitment of the AP1 transcription factor complex^68^, which mediates FGF signalling. The FGF pathway is critical for placode progenitor induction^69,70^ and Foxk2 may therefore participate in the activation of FGF responsive enhancers.

### scACE – a bioinformatics pipeline to allow accurate documentation and data accessibility

The analysis of complex data sets and integration of different data modalities requires a workflow of diverse bioinformatics tools. This is a fast-moving field with algorithms constantly being updated, making it challenging to provide transparent methods and analysis pipelines that accurately reproduce results presented. Furthermore, the lack of suitable documentation often prevents the community from exploiting valuable data sets. Here we present a Nextflow pipeline, scACE, for the analysis and integration of scATAC-seq and scRNA-seq, the generation of metacells and the identification of CREs with coordinated activities – so called accessibility modules. Finally, our pipeline implements a GRN inference approach which allows the prediction of new key nodes. scACE is user-friendly, accessible, and produces highly reproducible results, which in turn is key for others to explore our data. In addition, we make our data accessible using a ShinyApp, which allows exploration and visualization of our data.

A novel feature of scACE is the calculation of accessibility modules – chromatin regions with coordinated dynamics. While gene module analysis identifies co-regulated genes, we have adapted this approach to identify groups of putative CREs that open and close together in an unbiased manner. Modelling the dynamics of a set of coordinated genes is more informative than assessing individual genes, while also allowing unbiased analysis and minimising the loss of potentially important genes^4,23^. Here we demonstrate that defining accessibility modules, has the potential to reveal new biological insight. It has not only allowed us to characterise global changes in the chromatin landscape but also reveals that neural crest cells share accessibility profiles with placodes and neural cells supporting the idea that they retain some level of multipotency even at later stages^9,51^.

## Methods

### Key resources table

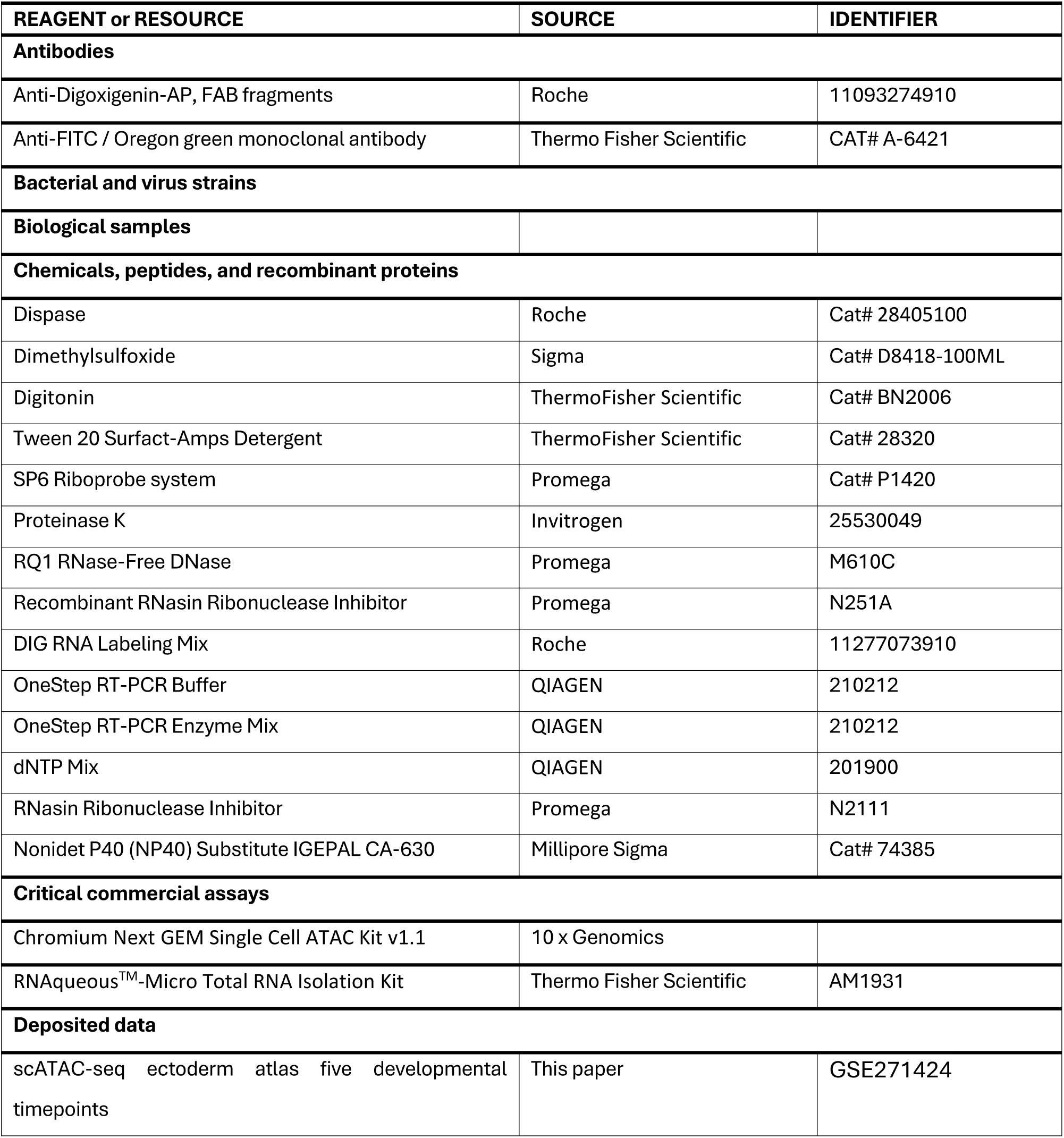
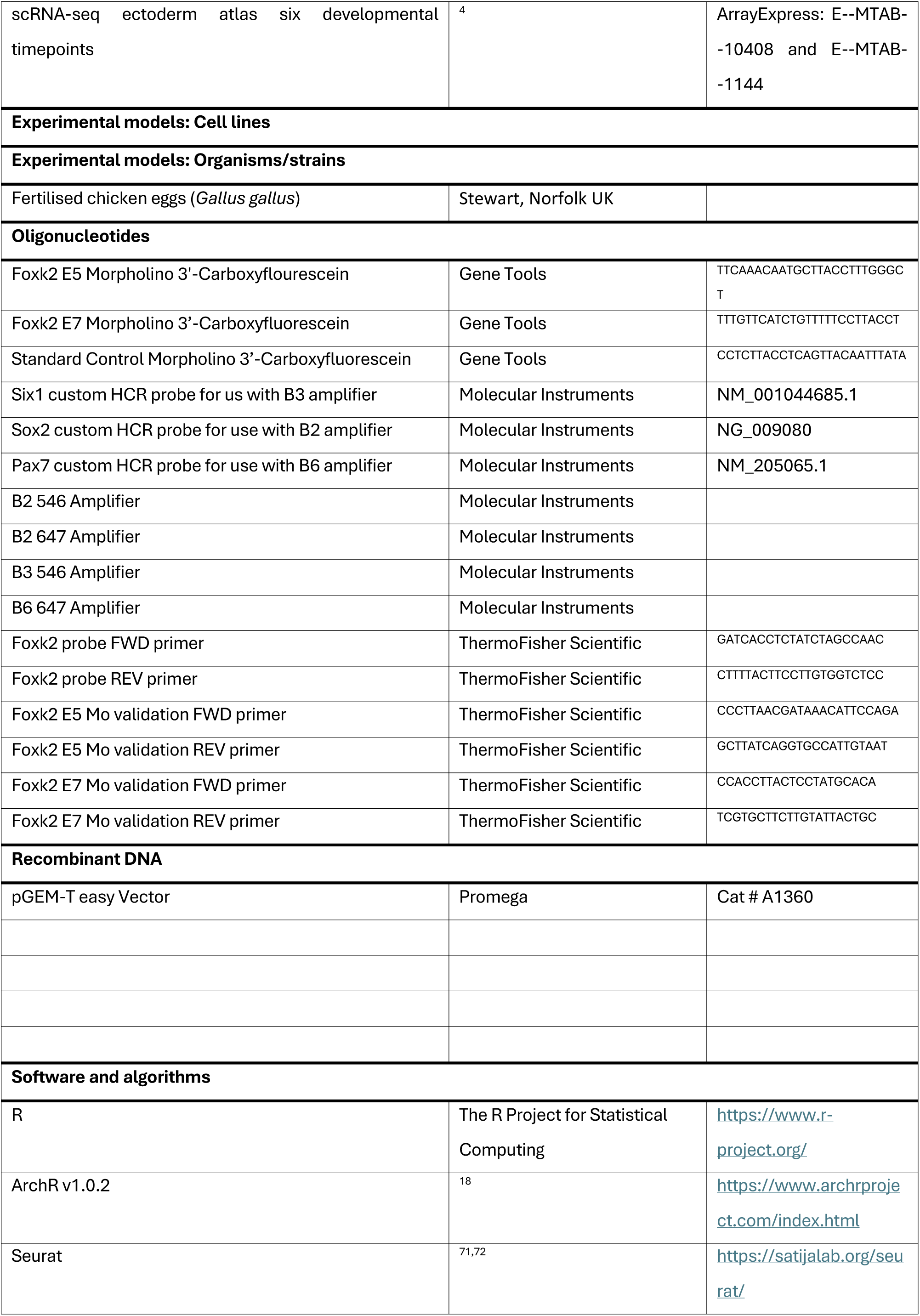
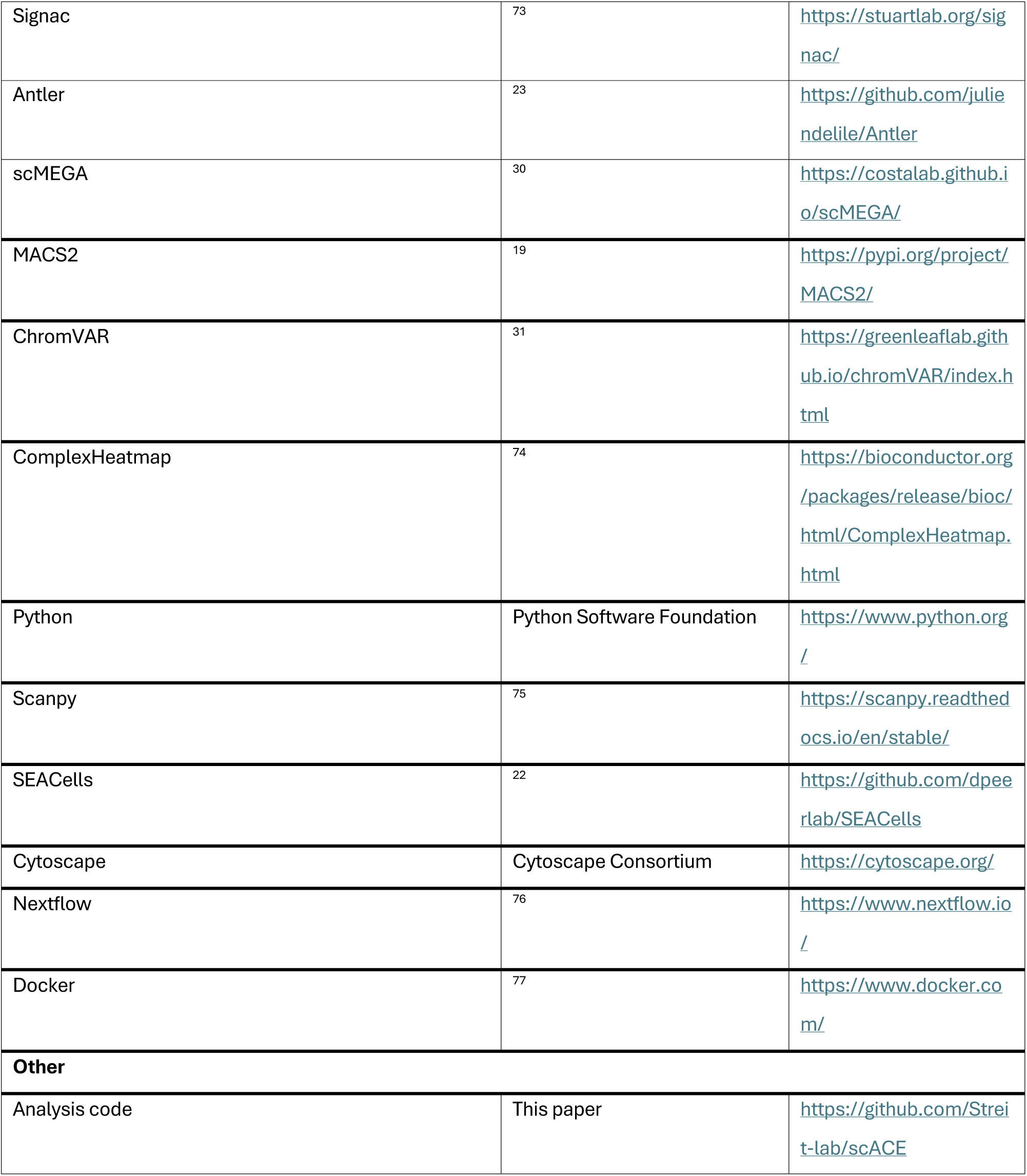

### Tissue collection

Fertilised chicken eggs (Stewart, Norfolk UK) were incubated at 38°C until they had reached Hamburger and Hamilton^17^ (HH) HH5, HH6, HH7, HH8^+^ (somite stage ss4) and HH9^+^ (ss8). Embryos were collected in Tyrode’s saline and ectoderm containing neural plate border cells was dissected manually using dispase (10 mg/ml) (Roche, 28405100). Tissues were cryopreserved at -80°C in 10% dimethyl sulfoxide (Sigma, D8418-100ML) in Tyrode’s saline until processing for single nuclei assay for transposase-accessible chromatin with sequencing (ATAC-seq).

### Nuclei isolation and scATAC sequencing

Nuclei isolation was carried out using a modified version of the 10x Genomics protocol (‘Nuclei Isolation from Mouse Brain Tissue for Single Cell ATAC Sequencing, Document Number CG000212 Rev B, 10x Genomic’, 2019). Samples were warmed in a 37°C water bath for 1 minute before adding 500 μl warmed Tyrode’s saline. Samples were spun at 4°C for 5 minutes at 200 relative centrifugal force (RCF), the supernatant was removed and replaced with 100 μl of 0.05X lysis buffer (see 10x protocol for composition). Samples were incubated on ice for 5 minutes before adding 1 ml wash buffer (see 10x protocol for composition). Nuclei were filtered (20 μm Miltenyi cell strainer) and recovered by spinning at 4°C at 500 RCF for 5 minutes. Nuclei were resuspended in nuclei solution buffer (10x Genomics, PN 2000153/2000207) and underwent single-cell ATAC-sequencing. Library preparation was carried out using the Chromium Next GEM Single Cell ATAC Kit v1.1 (10 x Genomics), before sequencing on the HiSeq 4000 100bp PE (Illumina) to a target depth of 50 k reads/cell.

### Data processing

Reads were aligned using 10x Genomics Cell Ranger-ATAC 2.0.0 against the chicken reference genome GRCg6a, using a modified GalGal6 ensembl 97 GTF file^4^. All scATAC-seq processing and visualisations were performed using the package ArchR^18^.

Potential doublets or clumps of nuclei were removed by filtering cells with a high number of fragments. The maximum fragment-counts threshold was defined as one standard deviation above the mean. This threshold was determined separately for each developmental stage to account for differences in sequencing depth between samples. Poor quality nuclei were removed using transcription start site (TSS) enrichment and Nucleosome Signal Ratio. Cells from each stage were clustered at a resolution of 3.6; cell clusters outside the 20-80% percentile range were removed. This approach was iterated until no more outlier clusters were found.

Dimensionality reduction was performed using the ArchR function ‘addIterativeLSI’^18^. Clustering was performed using the Shared Nearest Neighbour (SNN) approach as implemented by the Seurat package^71,78^. A clustering resolution of 0.8 was set for the full dataset and 0.7 for each individual stage, based on inspection of cluster tree plots which indicated that higher resolution caused many cells to switch cluster identities.

Peak calling was performed using ArchR’s function ‘addReproduciblePeakSet’^18^. First, two pseudoreplicates were generated for each cluster and stage using ArchR’s function ‘addGroupCoverages’^18^. Next, peaks were called on each pseudoreplicate using the MACS2 package ^19^ and ArchR’s iterative overlap method was used to create a consensus set of reproducible peaks.

To determine differentially accessible regions between clusters of cells we used the ArchR function ‘getMarkerFeatures()’. Differentially accessible peaks were visualised using the ArchR’s implementation of heatmap ‘markerHeatmap’^18^. Volcano plots were generated using custom R code.

### Cell state annotation

Previously published scRNA-seq data from chick neural plate border (NPB) cells at the same chick stage^4^ were used to label scATAC-seq cells with appropriate cell states. Label transfer was performed independently on each stage. The ArchR function ‘addGeneIntegrationMatrix’ was used to identify most similar cells between the both datasets and to transfer information, including cell state labels and gene expression information, from the scRNA-seq to the scATAC-seq data^18^.

### Metacell analysis and processing

Metacell analysis was performed using the SEACells package^22^; Seurat^71^ and SCANPY^75^ were used to handle the single cell datasets. Metacells were generated for both scRNA-seq and scATAC-seq data independently for each stage. The average SEACell size parameter was set to 10 for scRNA-seq data and 40 for scATAC-seq data. SEACells were created based on the previously calculated dimensionality reductions PCA for scRNA-seq and LSI for scATAC-seq. Metacells were initialized with 10 eigenvalues; the model was then fitted iteratively using default parameters with between 10 and 100 iterations. Hard assignments were used, soft assignments were visualised to check that generally cells belong to only one metacell. After generation, metacells were assessed for compaction, separation and cell state purity using functions from the SEACells package.

Both RNA and ATAC metacells were processed using the standard Seurat pipeline in which metacell count data were normalised and scaled, and PCA was performed using the number of principle components determined by elbow plots. Batch effects were regressed out for the RNA metacells. Metacells were then clustered at a cluster resolution of 5 and annotated using a previously published automated annotation pipeline^4^.

RNA and ATAC metacells were integrated using the SEACells tutorial^22^ exploring k values between 1 and 20. This process identified multiple pairs between single RNA and ATAC cells. A custom R script was created to unify these multiple pairs into one consensus annotation so that each single ATAC cell was labelled with a single cell state. This was achieved by substituting broad cell-state categories when an ATAC cell was assigned two similar labels (e.g. if an ATAC cell was labelled as both NC and dNC, it would be annotated as NC).

### Accessibility module identification and analysis

The aggregated peak count matrix across metacells was used as an input for accessibility module analysis to identify peaks that open and close together. Aggregated counts were normalised by conversion of raw counts to counts per 10,000. Peaks in promoter and exonic regions were removed. Peaks with the highest variability (top10,000 by variance) were retained and their count metacell matrix was processed using the Antler package to generate accessibility modules^23^. In short, an Antler object was created and counts were normalised to counts per million (CPM). Then, modules were identified with the parameters: corr_min = 3, mod_min_cell = 5, mod_consistency_thres = 0.3. The corr_t parameter was varied depending on the input data (0.3 for the full data, 0.3 for HH5, 0.4 for HH6, 0.6 for HH7, ss4 and ss8.

Motif enrichment analysis for peaks in each accessibility module was performed using function HOMER’s ‘findMotifsGenome.pl’ function^79^. A custom Nextflow^76^ module and bash script was created to take a list of peaks such as accessibility modules and transform them into a bed file which can be used as input. The ‘knownResults.html’ output file was inspected to identify known motifs with a q-value of < 0.05.

Target genes of peaks within each accessibility module were identified as the nearest gene on the genome, this was calculated at the time of peak calling in the ArchR object and was accessed via the ArchR function ‘getPeakSet’^18^.

Accessibility modules were visualised on a heatmap, as feature plots and as GAM plots. Accessibility module heatmaps were made using custom R functions that utilise the ComplexHeatmap^74^ R package. Feature plots were made using Seurat^71^. GAM plots were made using previously published modified custom R functions^4^.

### GRN inference

GRN inference was performed using the ArchR and scMEGA packages. The GRN was inferred on a placodal trajectory based on previously calculated latent time and lineage absorption probabilities^4^. Previously published lineage absorption probabilities^4^ were annotated on scATAC-seq data using the cell state annotation process described above. Cells with a placodal lineage absorption probability below 0.25 were not considered in the placodal trajectory.

Genes were associated to predicted enhancers using using ArchR’s function ‘addPeak2GeneLinks’^18^ with default parameters apart from: maxDist = 250,000, corCutoff = 0, FDRCutOff = 0.01.

Motif annotations were added to peaks in the scATAC data using the ArchR function ‘addMotifAnnotations’^18^. Motifs were obtained from the JASPAR 2020database collection ‘core’ taxonomy grouping ‘vertebrates’. Once motifs were annotated in the data, ArchR implementation of ‘ChromVar’^31^ (with default parameters) was used to calculate bias-corrected accessibility deviations for each cell. These deviation scores were used as a proxy for TF activity.

The R package scMEGA^30^ was then used to infer the GRN underlying placodal lineage segregation. TFs were selected by correlating TF activity with its expression using threshold parameters (cor.cutoff = 0.3, p.cutoff = 0.01). Genes were selected by variation across the trajectory (var.cutoff.gene = 0.3) and correlation with a linked peak (cor.cutoff = 0.3, fdr.cutoff = 1e-04). Peaks had to be a minimum of 2kb and maximum of 250kb away from the linked gene’s TSS. scMEGA vizualition across trajectories was performed using the scMEGA plotting functions using default parameters. Predicted interactions were filtered based on the correlation between TF activity and target gene expression (fdr < 0.01).

### Network visualization

A subnetwork of the GRN was visualised using Cytoscape software. Known placodal TFs and Foxk2 were visualised, and nodes coloured based on their temporal assignment calculated using scMEGA^30^.

### Chick embryo culture and electroporation

Fertilized hens’ eggs were incubated at 38 °C for 17 hours to reach stage HH4^17^. Splice modifying Foxk2 morpholinos targeting exons 5 and 7 were mixed and injected unilaterally at a total concentration of 1.5 mM (0.75 mM each), while a single control morpholino was injected at 1.5 mM. Injection mixes included fast green FCS dye, 3.6 % glucose, and 0.3 μg/μl carrier plasmid DNA. Electroporations were performed with the Ovodyne electroporator (TSS20, Intracel), using five 50 ms pulses of 5 V at 750 ms intervals^80^.

Embryos were collected using filter paper rings^81^ and cultured on egg albumin for a further 12 hours after electroporation, or until embryos reached stage HH7 to HH8^17^. Embryos were screened for FITC using an Olympus SZX12 fluorescent microscope to ensure successful uptake of morpholinos. Embryos were then fixed in 4% PFA for 1 hour at room temperature before being dehydrated in a series of methanol in PBT washes and stored at -20 °C.

### Whole mount *in situ* hybridisation, hybridisation chain reaction (HCR) and immunofluorescence (IF)

Colorimetric *in situ* hybridisation was performed following previously described protocols^82^. A DIG-labelled antisense RNA probe for Foxk2 was synthesised using an SP6-mediated transcription reaction with a linearised pGEM®-T easy vector containing a 297 bp Foxk2 amplicon. Pictures were taken with a Retiga2000R camera fitted to an Olympus SZX12 fitted and Q-Capture Pro7 software.

HCR v3 was carried out using the Molecular Instruments protocol^83^ adapted for whole mount chick embryos as described previously^84^. After rehydration, embryos were permeabilised by proteinase-K treatment (20 mg/mL; 2 mins) and postfixed in 4% PFA for 20 mins. After post-amplification washes, embryos were incubated for 10 mins in DAPI (10 mg/mL).

Immunofluorescence using an anti-FITC antibody conjugated with Alexa Fluor™ 488 was performed after the HCR to visualise the morpholino against Six1 expression. Embryos were washed in PBS-Tx (PBS + 0.2% Triton X-100) and blocked in 5% goat serum in PBS-Tx for 3 hours at room temperature, after which they were then incubated at 4 °C overnight in primary antibody diluted in blocking solution. Embryos were then washed five times for 30 minutes in PBS-Tx before being moved to 50% glycerol in PBS for imaging.

HCR IF embryos were imaged using the Zeiss LSM 980 confocal microscope with the 10X objective and Zen blue software. A z-stack was taken through each embryo with a slice interval of 8 μm. ImageJ was used to create maximum intensity z-projections and extract average pixel intensity values across electroporated regions of the embryo using the ROI manager tool. The ggplot2 package in R was used to plot ROI intensity values.

### Reverse Transcription PCR Morpholino validation

Following the electroporations described above, some embryos were chosen for validating the efficacy of Foxk2 morpholinos and thus their ability to induce a functional knockdown. Electroporated tissues of Foxk2 and control embryos were dissected and processed for tRNA extractions using the RNAqueous^TM^-Micro total RNA isolation kit (Invitrogen #AM1931), including DNase I treatment and DNase inactivation steps. 2 μL of tRNA product was subsequently used for a 20 μL volume OneStep RT-PCR (QIAGEN #210212) reaction with primer pairs flanking either Foxk2 exon 5 or exon 7. The thermocycler program included an initial 30 minutes at 50 °C for reverse transcription, followed by a touch down PCR of 12 cycles with annealing temperatures from 64 °C to 52 °C and a subsequent 22 cycles maintained at 52 °C. Denaturing and extension temperatures were set at 95 °C and 72 °C for 30 and 45 seconds, respectively, for each cycle, with a final extension time of 15 minutes. Finally, RT-PCR products were run on a gel to confirm the presence of Foxk2 transcripts missing exons 5 and 7 in response to the splice modifying Foxk2 morpholinos (Supplementary Fig. 11).

## Data and code availability

All raw and processed scATAC-seq data have been uploaded to GEO under GSE271424. The predicted regulatory interactions of the GRN are available in supplementary table 2. The code used for this study has been organized into a reproducible Nextflow pipeline and deposited on Github. It is publicly available at https://github.com/Streit-lab/scACE. Each process of the Nextflow pipeline utilizes Docker environment containers to ensure consistent software environments, these containers are publicly available on Dockerhub at https://hub.docker.com/u/alexthiery Any additional information required to reanalyze the data reported in this work paper is available from the lead contact upon request.

## Acknowledgements

The authors are grateful to Chantal Hubens and Robert Hayward for excellent technical support and to Claudio D. Stern and David Pearton for critical reading of the manuscript. This work was supported by the Wellcome Trust (108874/B/15/Z), the BBSRC (BB/S005536/1; BB/R006342/1) and in part by the Francis Crick Institute which receives its core funding from Cancer Research UK (FC001051), the UK Medical Research Council (FC001051), and the Wellcome Trust (FC001051).

## Author contributions

The project was conceived by AS, APT and EH; AS, APT and EH designed and interpreted experiments with contribution from ALB. EH together with APT designed the bioinformatics analysis pipeline with advice from AV. EH carried out all bioinformatics analysis; JL performed all functional analysis and together with EH implemented the ShinyApp. JB advised data analysis. AS and JB obtained funding.

## Declaration of Interest

The authors declare no competing interests.

## Notes

### Competing Interest Statement

The authors have declared no competing interest.

