## Supplementary Figures for "A single cell ATAC-seq atlas uncovers dynamic changes in chromatin accessibility during cell fate specification at the neural plate border"

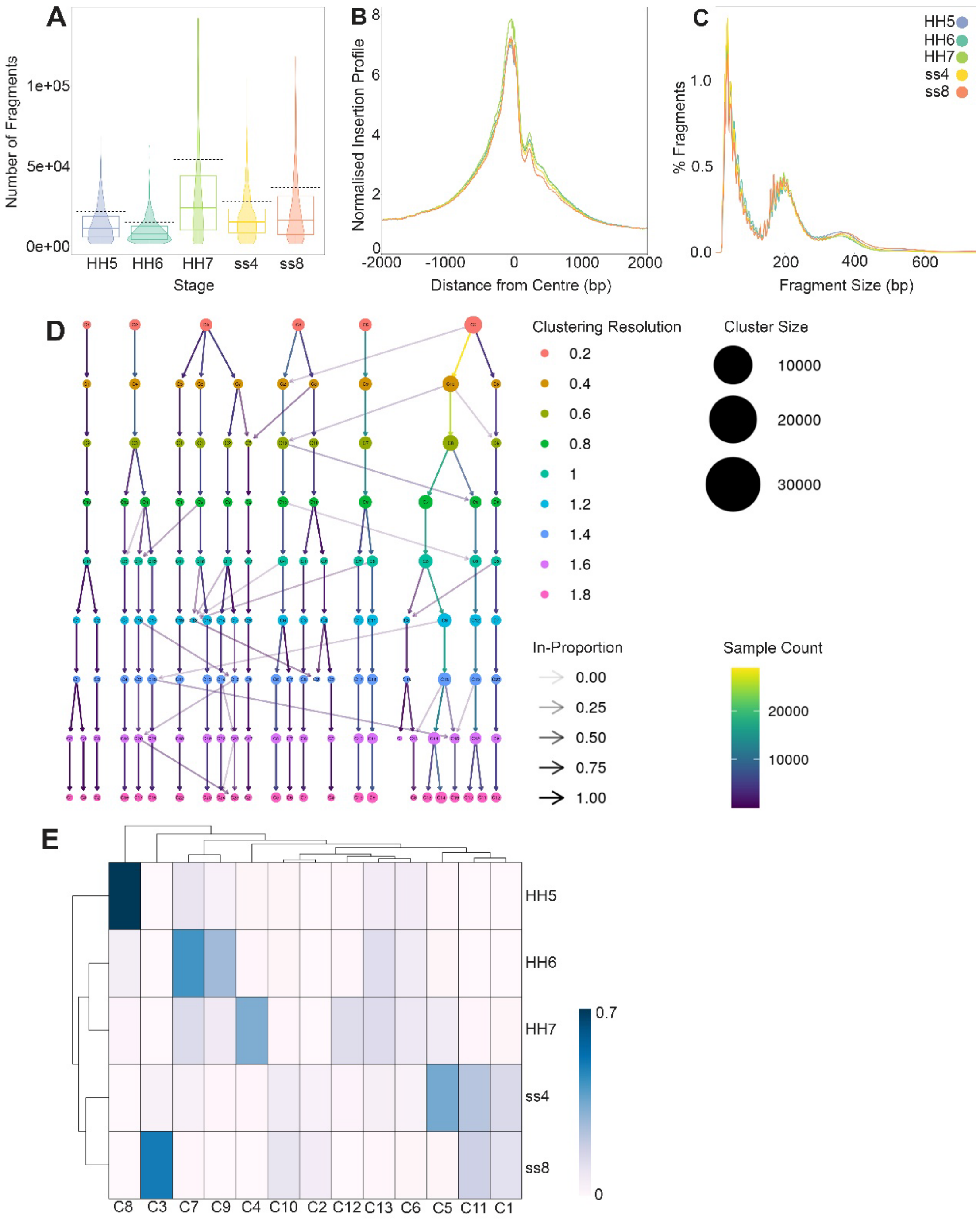


**Supplementary Figure 1. scACE quality control and filtering.** A. Violin plots showing distribution of fragment counts per sample. Dashed black lines indicate sample-centric thresholds for filtering (1SD above sample mean) B. Fragment size distribution plot showing expected nucleosome banding pattern across all samples. C. Transcription start site enrichment plot showing increased insertions around transcription start sites across all samples. D. Clustering tree showing proportion of cells falling into each cluster at incremental clustering resolutions from 0.2 to 1.8. E. Heatmap showing proportions of cells in each cluster coming from each sample.

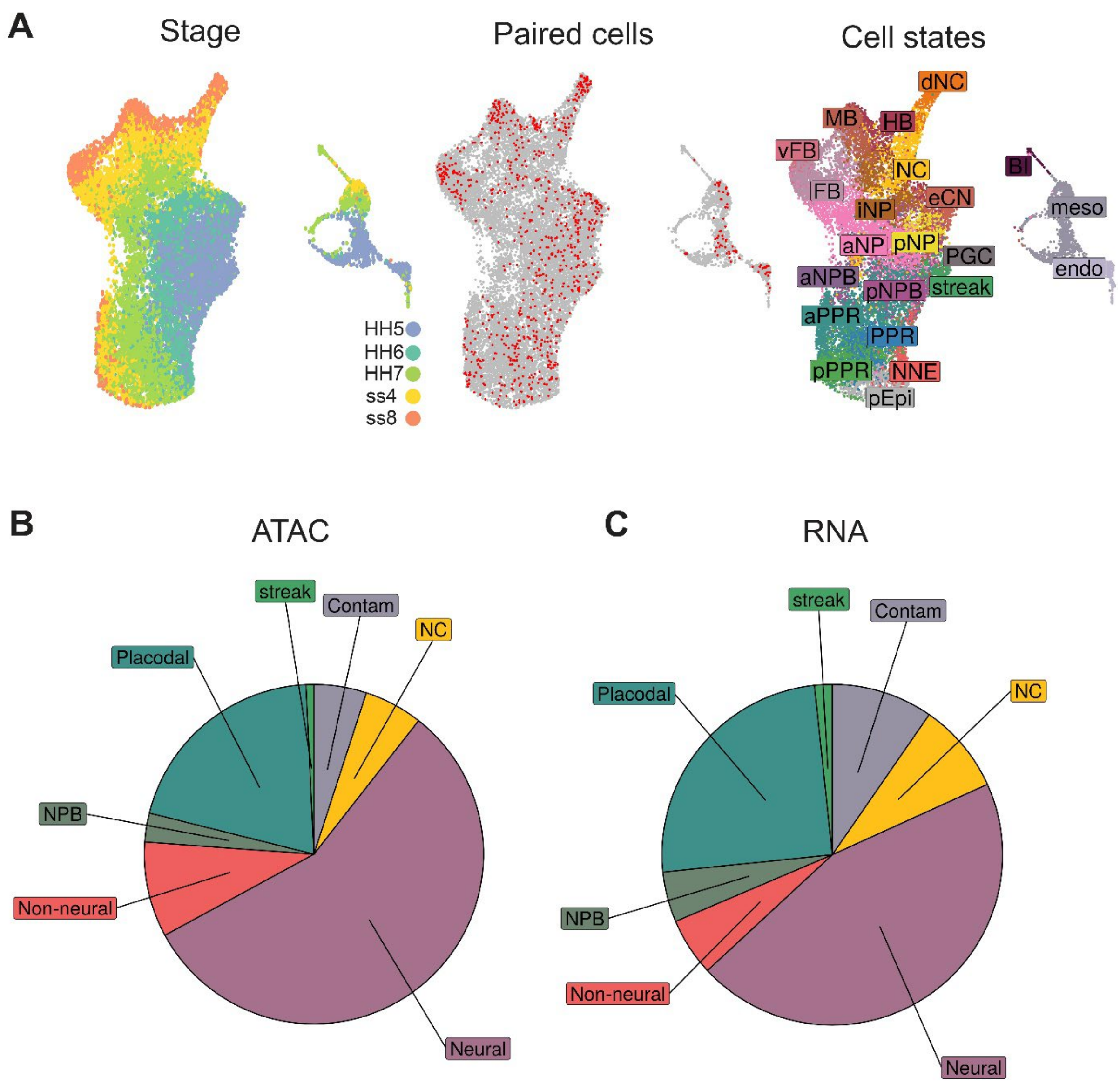


**Supplementary Figure 2 – scACE cross-modality integration and label transfer.** A. UMAP embeddings of scRNA-seq data collected from the chick ectoderm (Thiery et al., 2023). From left to right: UMAP showing developmental stage, UMAP showing distribution of scRNA-seq cells used for label transfer in red, cells not used for label transfer in grey, UMAP showing cell states. B. Piechart showing distribution of broad cell states in scATAC-seq data after label transfer. C. Piechart showing distribution of broad cell states in scRNA-seq data.

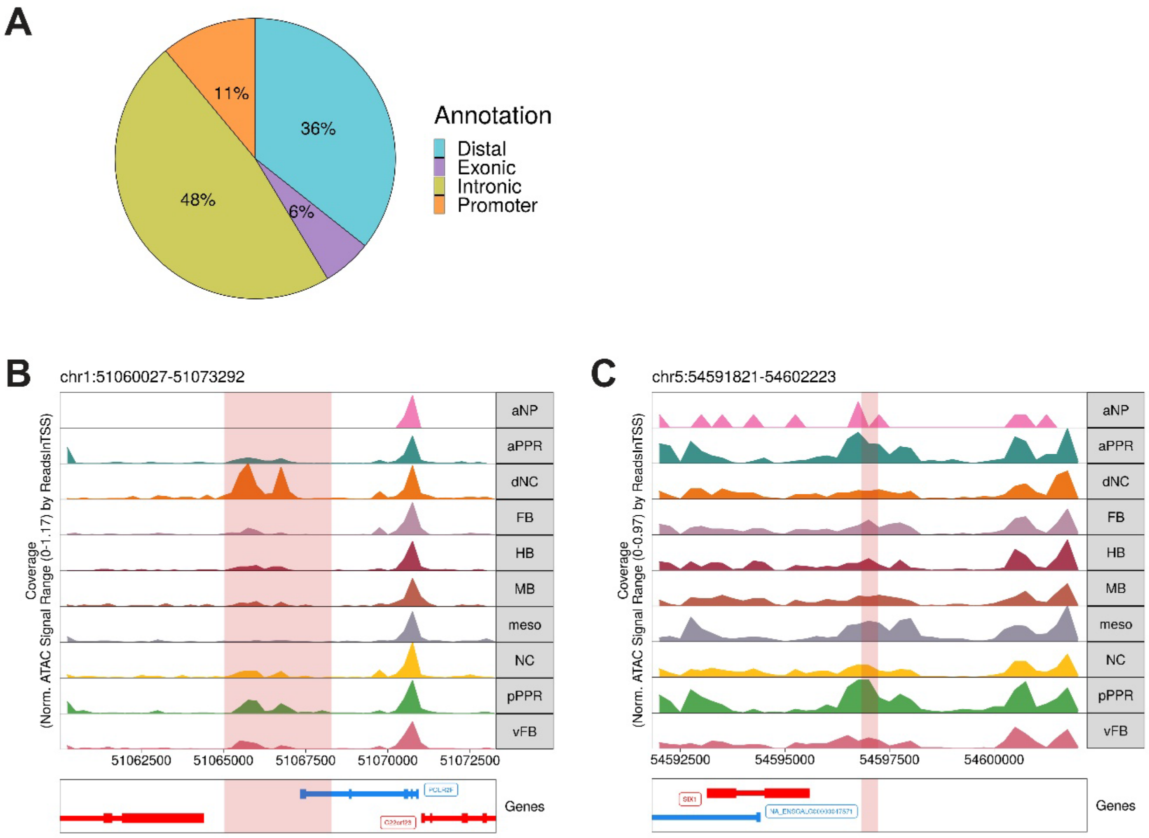


**Supplementary Figure 3. scACE peak calling and cross-reference to known enhancers.** A. Piechart showing percentage of peaks annotated as distal, exonic, intronic and promoter based on their genomic location. B. Genome browser plot showing accessibility profiles at ss8 aggregated over cells partitioned by their cell state centred around the location of the known enhancer Sox10_E1 (Betancur et al., 2010). Enhancer region is highlighted in red and browser plot shows 5,000bp on either side. C. Genome browser plot showing accessibility profiles at ss8 aggregated over cells partitioned by their cell state centred around location of known enhancer mSix1-21 (Sato et al., 2012); the plot shows the enhancer in red and 5,000bp flanking regions.

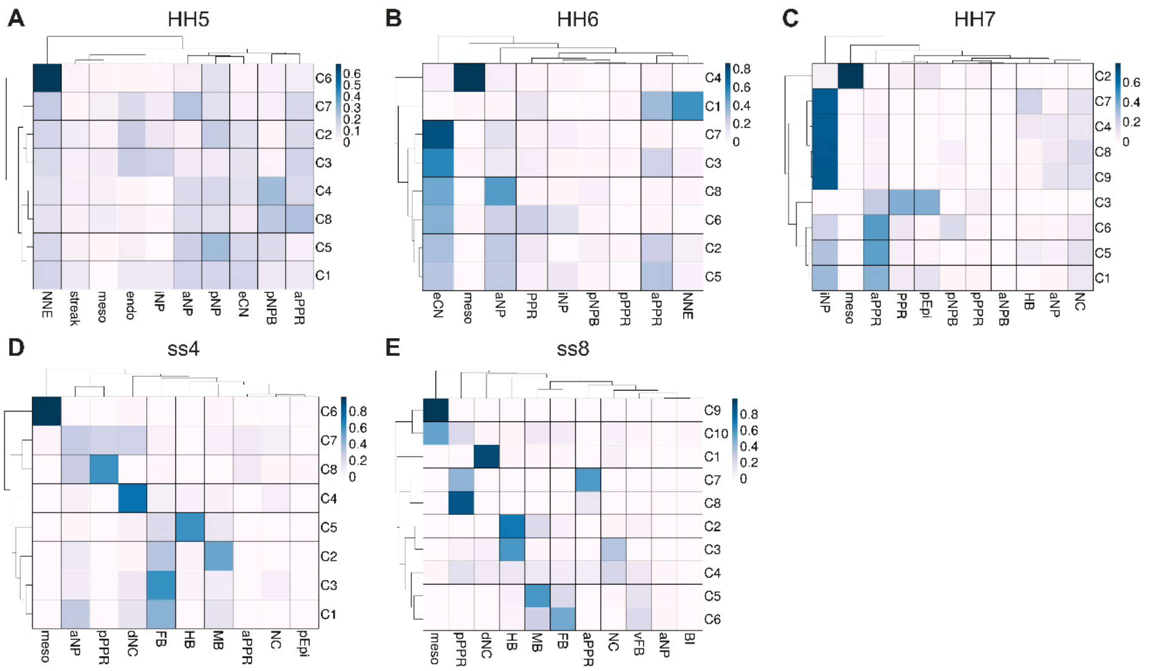


**Supplementary Figure 4. Distribution of cell state labels across clusters at each stage.** A-E Heatmap of each stage (HH5 to ss8) showing cell state labels across the x axis and cluster identities across the y axis. Colours indicate the proportion of cells (from 0 to 1) in each cluster that have been assigned the cell state label.

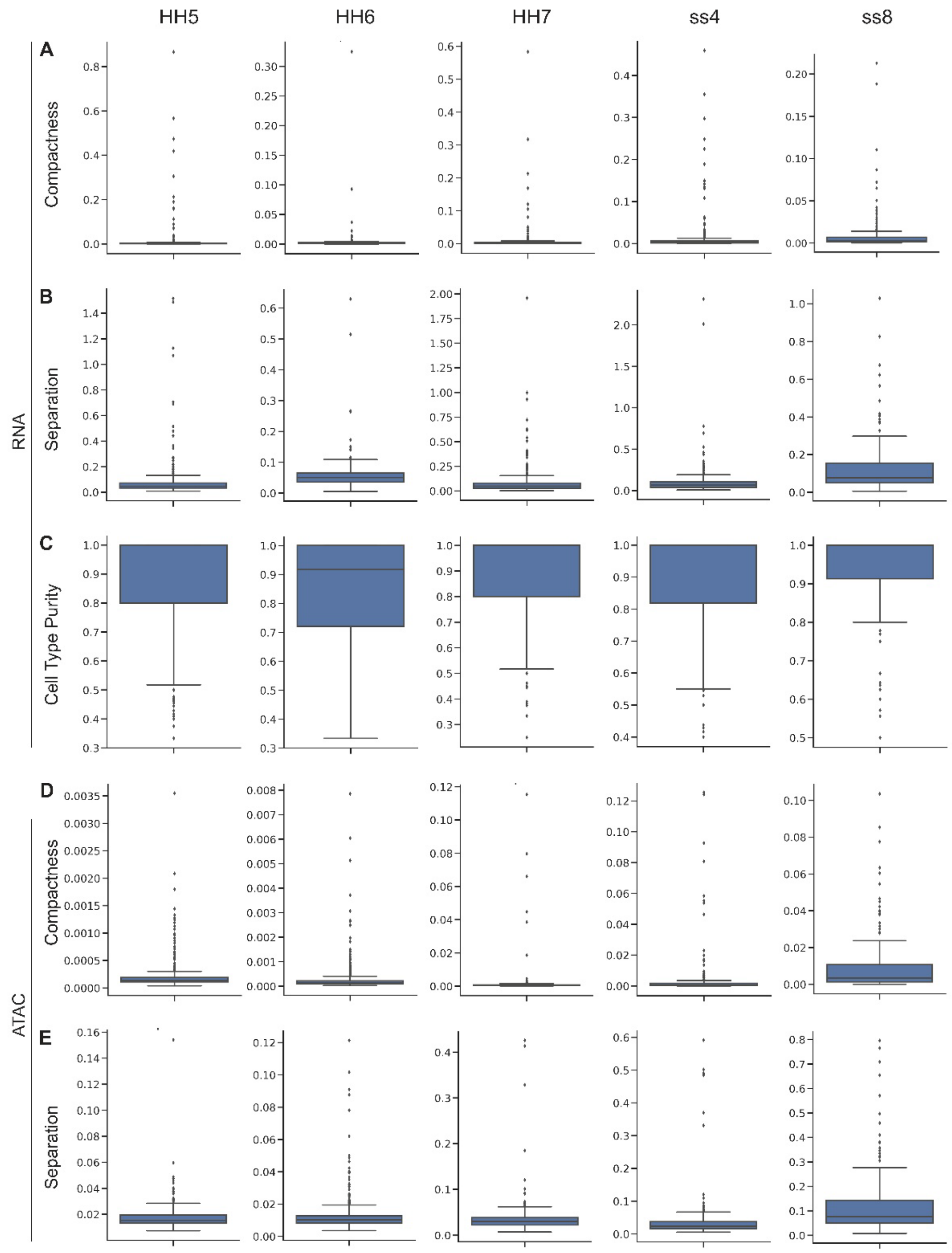


**Supplementary Figure 5. scACE metacell calculation and quality control.** A. Boxplots showing distribution of metacell scores for compactness across scRNA-seq samples. A low compactness score indicates that cells within a metacell are similar to each other. B. Boxplots showing distribution of metacell scores for separation across scRNA-seq samples. Higher separation scores indicate that metacells are more different from each other. C. Boxplots showing distribution of cell type purity scores across scRNA-seq samples. Metacells with a purity score close to 1 are made of cells from only one annotated cell type. D. Boxplots showing distribution of metacell scores for compactness across scATAC-seq samples. E. Boxplots showing distribution of metacell scores for separation across scATAC-seq samples.

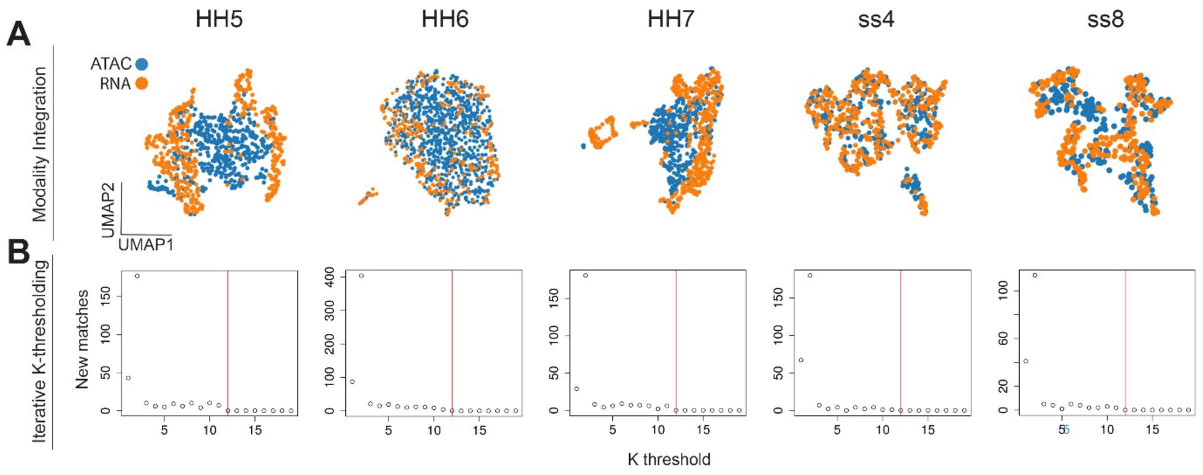


**Supplementary Figure 6. scACE metacell integration and label transfer.**

A. UMAPs coloured by modality to visualise projection of ATAC and RNA metacells onto a shared low dimensional space for each stage. B. Scatter plots showing number of additional new labels added with each increase of k values from 1 to 9. Red vertical line indicates the k threshold used for downstream analysis (k=12).

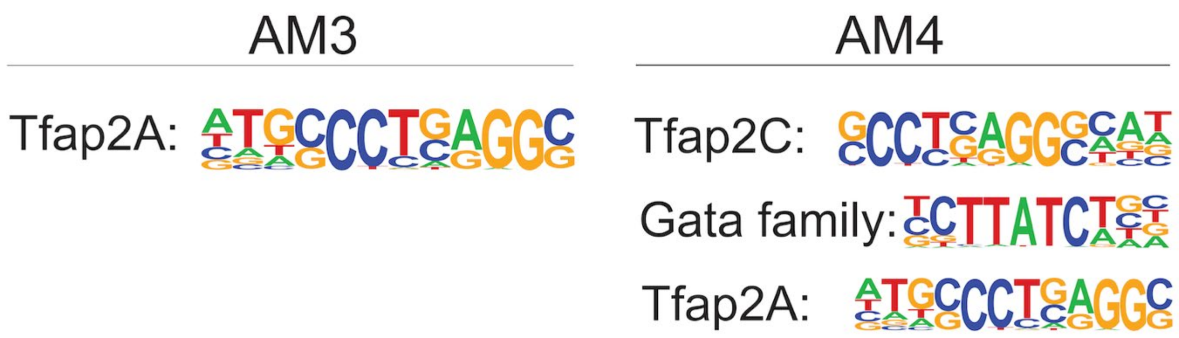


**Supplementary Figure 7. Motif enrichment in the placodal accessibility modules AM3 and AM4.**

Homer motif enrichment results for chromatin regions in AM3 (left) and AM4 (right) q-value < 0.05.

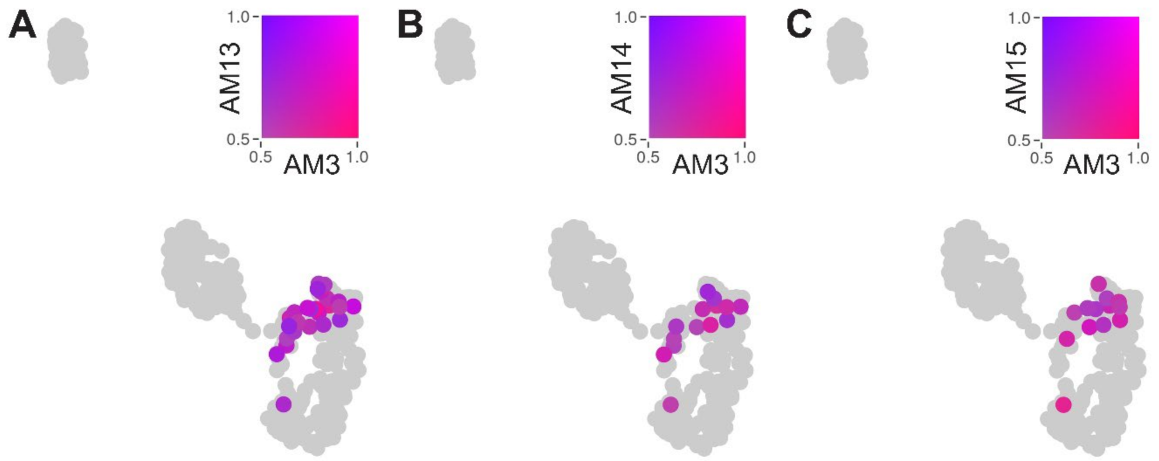


**Supplementary Figure 8. Co-accessibility of placodal AM3 and neural AMs in neural crest cells.** A. Feature plot showing co-accessibility of the neural AM13 and the placodal AM3 at ss4. B. Co-accessibility plot for neural AM14 and placodal AM3. C. Co-accessibility plot for neural AM15 and placodal AM3.

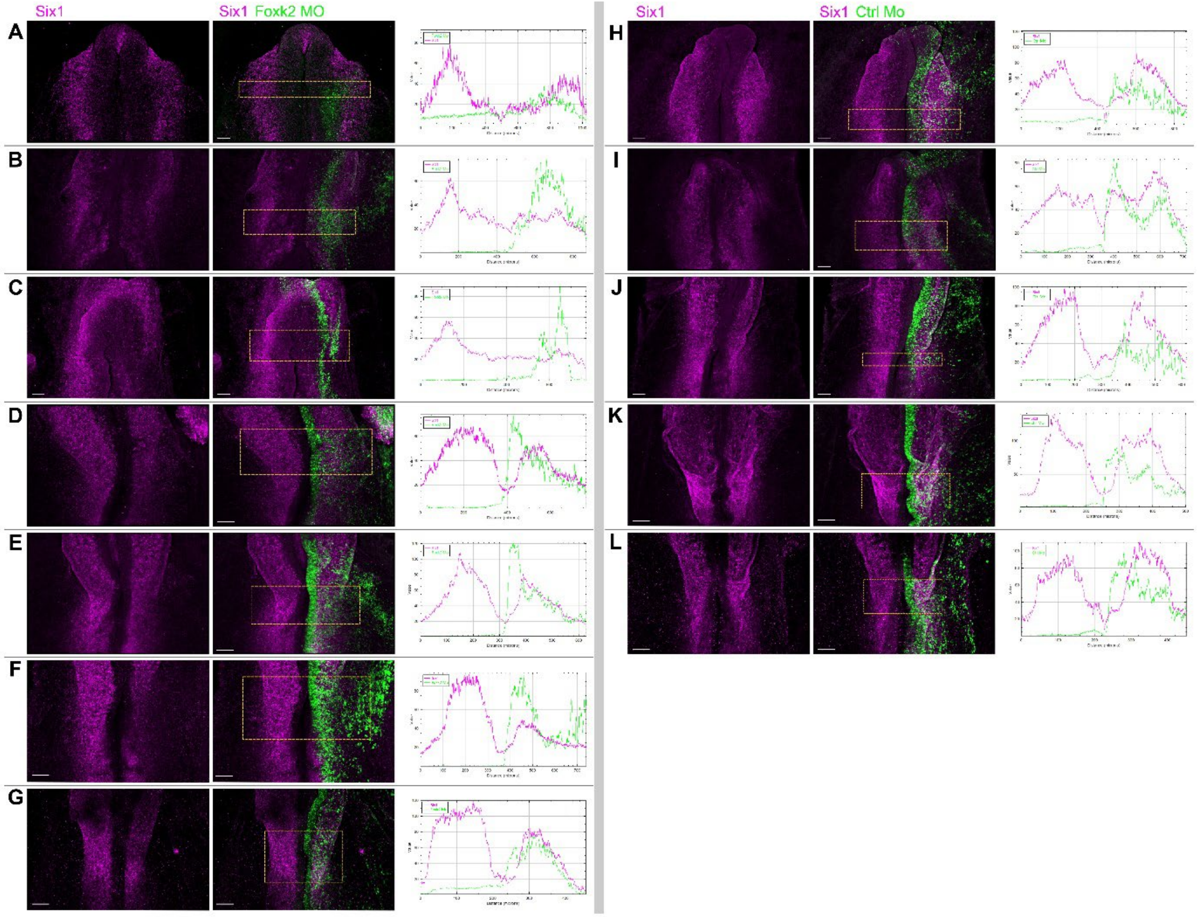


**Supplementary Figure 9. Foxk2 knock-down leads to reduction of the placodal marker *Six1*.** A-G. HH3-HH4 embryos were electroporated unilaterally with FoxK2 MOs (green), harvested at ss1-5 and processed for HCR in situ hybridisation for *Six1* (magenta). H-I. HH6 embryos were electroporated unilaterally with control MOs (green), harvested at ss1-5 and processed for HCR in situ hybridisation for *Six1* (magenta). Average fluorescent intensities for morpholino and *Six1* channels were quantified along the medio-lateral axis of the boxed areas and plotted to the side of each embryo.

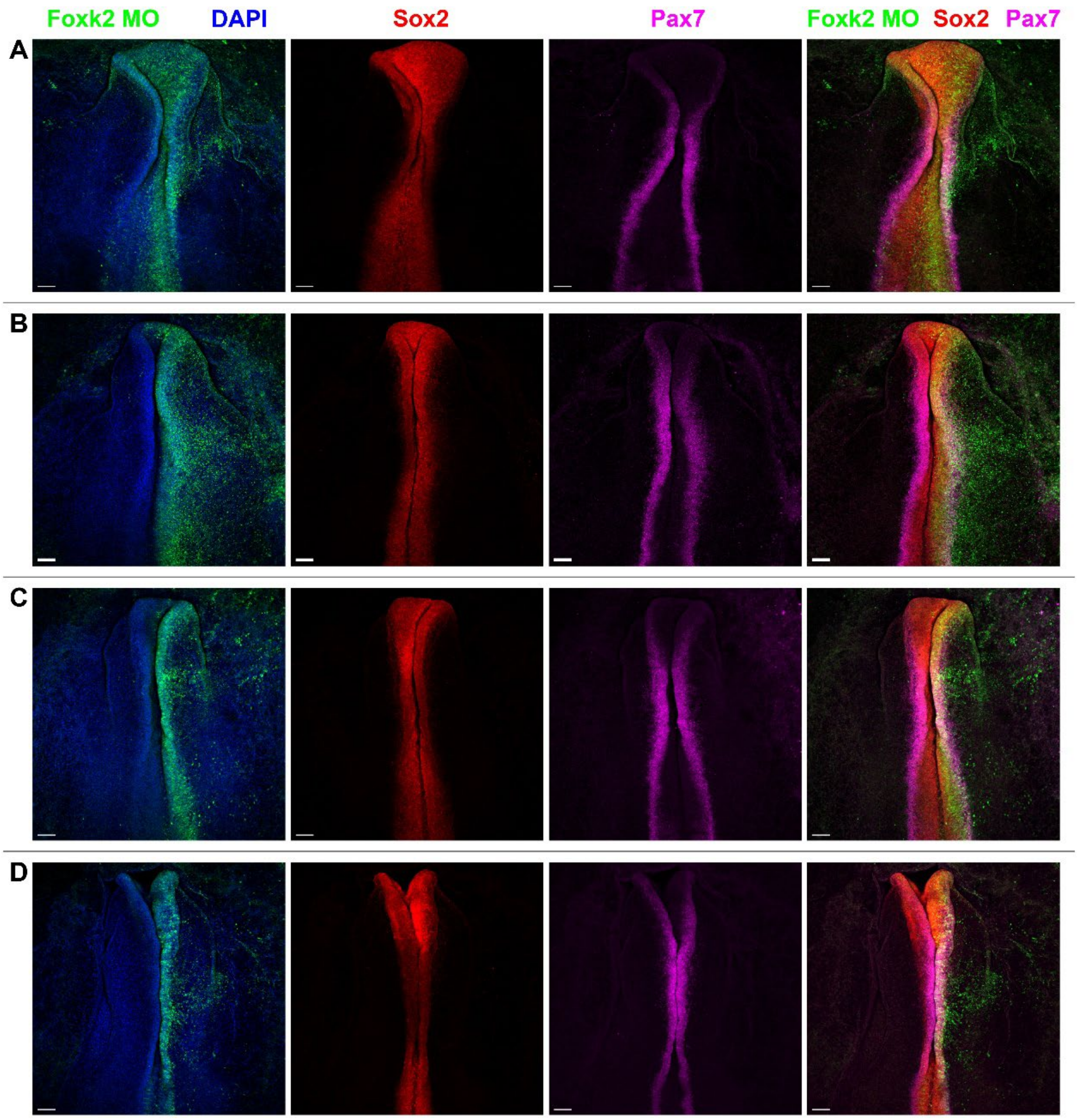


**Supplementary Figure 10. Foxk2 knock-down does not affect neural or neural crest marker expression.** A-D. HH3-4 stage embryos were unilaterally electroporated with Foxk2 splice modifying morpholinos. Embryos were harvested at ss1-5 and processed for HCR *in situ* hybridisation for the neural marker *Sox2* and the neural crest marker *Pax7*. *Sox2* (n=8) and *Pax7* (n=10) show normal expression patterns.

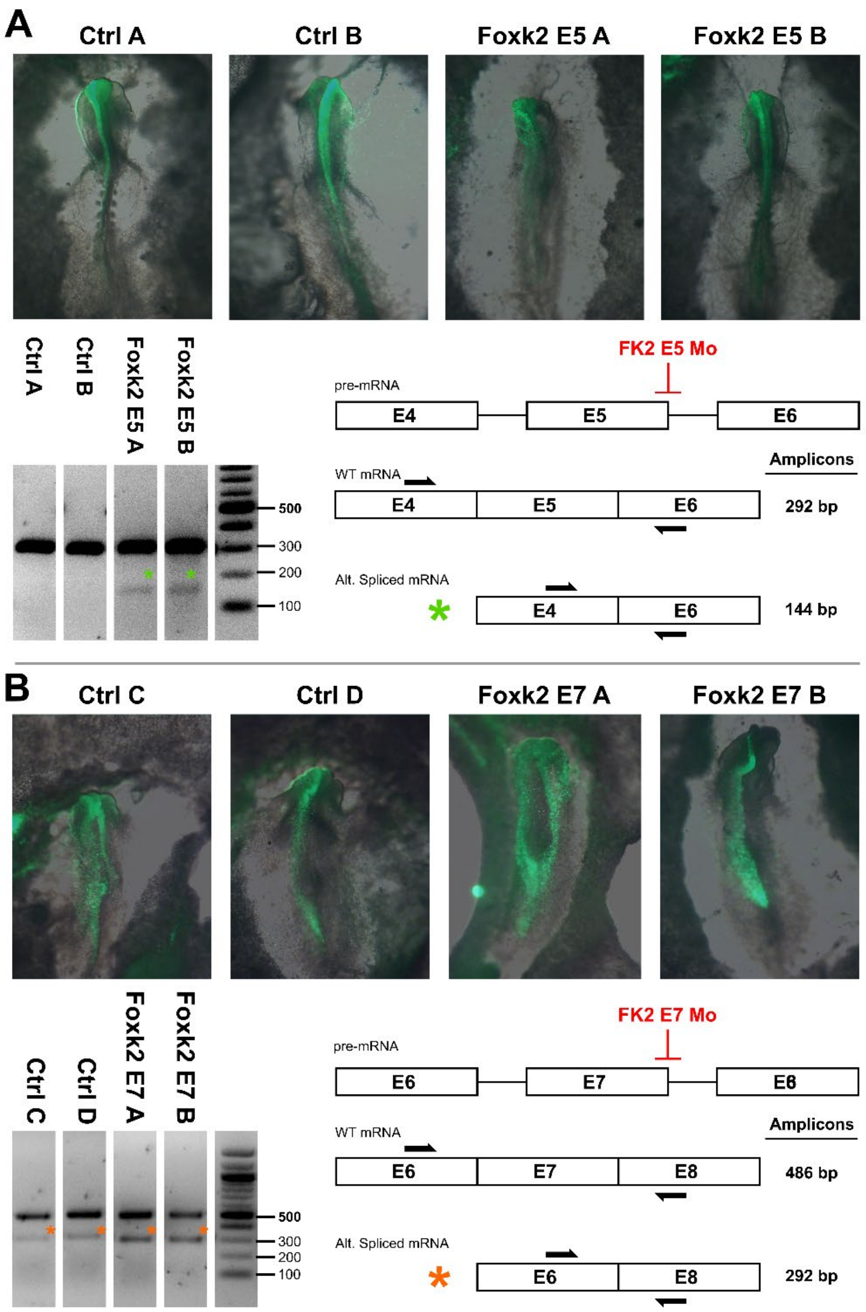


**Supplementary Figure 11. RT PCR validation of Foxk2 splice modifying morpholinos.** Foxk2 and control MO electroporated embryos were processed for RT PCR to assess the size of Foxk2 transcripts. A. Electroporations with a MO targeting Foxk2 exon 5 leads to exon deletion; electroporation is mosaic, and amplicons contain both the wildtype and exon 5 deleted bands (green asterisk). Control MO electroporated embryos show the wildtype band only. B. Electroporations with a MO targeting Foxk2 exon 7 also show both wildtype and exon 7 deleted bands (orange asterisk). However, controls show the same pattern albeit with lower levels of the smaller band suggesting the presence of a previously undescribed endogenous splice variant that lacks exon 7.
